# FTO-dependent m6A RNA dysregulation underlies memory deficits induced by early-life stress

**DOI:** 10.64898/2026.03.30.715262

**Authors:** Dipanjana Banerjee, Qiongyi Zhao, Sania Sultana, Sarbani Sammdar, Timothy Bredy, Sourav Banerjee

## Abstract

Cognitive functions in adults are mainly attributed to experience-dependent plasticity. Nonetheless, the developmental encoding of memory deficits is still inadequately addressed. Here, we demonstrate that early-life stress (ELS) reprograms the hippocampal epitranscriptome by enhancing N6-methyladenosine (m^6^A) deposits during early development leading to memory deficit in adulthood. We observed a shift toward hypermethylation of transcripts including coding and non-coding RNAs (lncRNAs) following maternal separation. We also observed that these transcripts encoding proteins necessary for translational regulation, ribosome biogenesis and mitochondrial function. This epitranscriptomic change is driven by ELS-induced downregulation of the m^6^A demethylase FTO (Fat mass and obesity-associated protein). We observe that the overexpression of FTO in young adult mice selectively rescues memory deficits without ameliorating elevated anxiety. Further, the knockdown of FTO in primary hippocampal neuron, mimicking ELS - induced reduction of its expression, leads to reduced translation as detected by puromycin labelling. Taken together, our study demonstrated previously uncharacterized mechanism of ELS-induced epitranscriptomic change linked with memory deficit via the regulation of protein synthesis.

## Introduction

m^6^A-RNA methylation (N6-methyladenosine) is the most abundant RNA modification in eukaryotes that fine-tunes an additional layer of RNA regulation (Akhtar et al., 2021; Sendinc & Shi, 2023); dynamically regulated by a set of functionally antagonistic enzymes called “writers” (methyltransferases) and “erasers” (demethylases) (Loedige et al., 2023). The writer complex comprises METTL3 (methyltransferase-like 3), METTL14 (Methyltransferase-like 14), and WTAP (Wilms tumor 1 associated protein) that catalyse the barcoding of RNA with m^6^A, while erasers like FTO (Fat mass and obesity-associated protein) and Alkbh5 (AlkB homolog 5) remove the modification from RNAs (Jia et al., 2011; Ping et al., 2014; P. Wang et al., 2016; Zheng et al., 2013). Another group of proteins called “readers,” e.g., YTHDF1-3 (YTH N6-methyladenosine RNA binding protein 1, 2, 3), recognise the m^6^A topography on the transcripts and channel them to specific signalling pathways (H. Shi et al., 2018; Zaccara & Jaffrey, 2020). While the role of m^6^A in RNA stability and translation has been widely studied (Zhou et al. 2018, Shi et al., 2018), its contribution to the regulation of early-life stress (ELS) remains elusive.

Previous studies have converged on the effects of ELS on long-term cognitive health, particularly in the form of heightened anxiety and memory deficits (Levin & Liu, 2021; Lin et al., 2016; Pattwell & Bath, 2017; A. Wang et al., 2020). The hippocampus, with its dual role in regulating emotion and spatial memory, has also been shown to be affected by ELS (Bertagna et al., 2021; Brosens et al., 2023; Hanson et al., 2015; Lee et al., 2019). Still, the molecular underpinnings of the hippocampal circuitry modulating such behavioural patterns remain poorly understood. Emerging evidence also suggests that RNA modifications, particularly m^6^A, play a critical role in regulating RNA stability and expression under stress conditions (Anders et al., 2018; Kisliouk et al., 2020). Here, we investigated how m6A-RNA modification influences hippocampal circuitry in the context of ELS using the long-standing paradigm of maternal separation (MS) (Sood et al., 2018).

Our study shows that maternal separation causes an increase in m^6^A-RNA deposition along the transcripts in the hippocampus of mice. The impact of change in m^6^A topography extends beyond protein-coding RNAs to include long non-coding RNAs (lncRNAs). LncRNAs are more than 200 nucleotides long and typically do not contain open reading frames for encoding proteins (Fatica & Bozzoni, 2014; Ma et al., 2013). This subset of RNAs has also been increasingly evidenced to have a role in translation and synaptic plasticity (Banerjee et al., 2024; Mattick et al., 2023; Mercer et al., 2008, 2009; C. Shi et al., 2017; A. Wang et al., 2017) and for their regulatory roles in the brain (Briggs et al., 2015; Ernst & Morton, 2013; Samaddar et al., 2023). The hypermethylation of hippocampal transcripts coincides with a significant downregulation of the RNA demethylase FTO, suggesting that the hippocampus is particularly vulnerable to m6A-mediated epigenetic shifts. Reduced FTO expression is directly correlated to the spatial memory deficits observed after ELS, and its overexpression rescues the phenotype. The downregulation of FTO in hippocampal neurons leads to a decrease in global protein synthesis. Collectively, our study reveals a molecular framework involving RNA methylation that may explain the cognitive deficits observed in adults who have been exposed to ELS.

## Materials and Methods

### Ethical clearance for animal maintenance

The animals used in this study were housed following the guidelines established by the Institutional Animal Ethics Committee of the National Brain Research Centre (NBRC-IAEC) in India. The study protocols received approval under reference numbers NBRC/IAEC/2015/172 and NBRC/IAEC/2020/178. NBRC-IAEC is accredited by the Committee for the Purpose of Control and Supervision of Experiments on Animals (CPCSEA) under the Ministry of Fisheries, Animal Husbandry, and Dairying, Government of India, with registration number 464/GO/ReBi-S/Re-L/01/CPCSEA. C57BL/6 and CD1 mice strains were housed individually in standard cages under a 12-hour light/dark cycle (6 a.m. to 6 p.m.). The mice had ad libitum access to food and water. Environmental conditions were maintained at a constant temperature of 26 ± 1°C and a humidity level of 40 ± 5%.

### Maternal Separation

To induce ELS, we explored the maternal separation paradigm as a potent tool (Alves et al., 2022). C57BL/6 pregnant dams were subject to random allocation, dividing them into either a control cohort or a maternally separated group. A litter size of 6-8 pups was maintained. The separation started from postnatal day 2 (P2) and extended until P14, with a daily separation period of 3 hours, from 0900 to 1200. Before each separation session, the mother was relocated to a fresh cage, while the pups were transferred to a different room. The pups were gently removed from their original cage individually and placed on soft cotton bedding within a glass beaker positioned on a heating pad, maintaining a consistent temperature range of 36°C to 38°C. The inner wall of the glass beaker was lined with paper to prevent singeing. Upon completion of the separation period, the pups were carefully returned to their original cage, followed by the reintroduction of the mother. Subsequently, the animals were kept under observation to ensure the mother’s engagement in tending and nursing the pups. The pregnant dams comprising the control group experienced minimal disruption, with cage changes occurring once every three days, consistent across all experimental conditions. At P21, male pups were selected for the harvesting of hippocampal tissue, facilitating further investigation and analysis.

### 2.3. Immunoprecipitation and dot-blot

To perform m^6^A RNA immunoprecipitation (MeRIP) assay, hippocampi were isolated from P21 C57BL/6 male mice and collected in an immunoprecipitation buffer containing 20 mM Tris-HCl (pH 7.5), 100 mM KCl, 5 mM MgCl2, 0.5% NP-40, 1 mM DTT, 0.5 µg/µl heparin, protease inhibitor (Sigma; 1 µl/ml), phosphatase inhibitor (Sigma; 1 µl/ml), and RNase inhibitor (SuperaseIn; 100 U/ml). The hippocampi were homogenised into a single-cell suspension. Cell debris was removed by centrifugation at 10,000g for 5 minutes at 4°C, and the supernatant was collected. 10% of the total crude lysate was kept aside as total input. Simultaneously, agarose beads were prepared by washing three times with wash buffer 1 (20 mM Tris-HCl, pH 7.5, 100 mM KCl, 5 mM MgCl2, 0.5% NP-40, 0.5 µg/µl heparin, and RNase inhibitor) at 10,000g for 5 minutes at 4°C. The beads were then incubated in 2.5% bovine serum albumin (BSA), prepared in wash buffer 1, for 1 hour at 4°C, 50 rpm. After incubation, the beads were precipitated by centrifugation at 10,000g for 5 minutes at 4°C, and the BSA solution was decanted. The collected supernatant was added to the BSA-incubated beads for pre-cleaning, incubating for 1 hour at 4°C, 50 rpm. The beads were then precipitated again at 10,000g for 5 minutes at 4°C, and the supernatant was collected and divided into two equal parts. Rabbit m6A antibody was added to one half, and rabbit IgG was added to the other half, followed by incubation for 4 hours at 4°C with gentle shaking. The RNA-antibody mixture was then added to agarose beads pre-equilibrated in wash buffer 1 (by washing three times at 10,000g for 5 minutes at 4°C) and incubated for 2 hours at 4°C, 50 rpm to allow the antibody-RNA complexes to bind to the beads. Following incubation, the beads were washed twice with a high-salt wash buffer (20 mM Tris-HCl, pH 7.5, 150 mM KCl, 5 mM MgCl2, 0.5% NP-40, and RNase inhibitor) and then twice with a normal wash buffer (20 mM Tris-HCl, pH 7.5, 100 mM KCl, 5 mM MgCl2, 0.5% NP-40, and RNase inhibitor) to remove non-specifically bound RNA. 1 µl of bead was used for dot-blot and the rest of the beads were stored at −80 °C.

1 µl of the sample from the total input, m6A-IP, and IgG-IP was dotted onto a nitrocellulose membrane. The membrane was incubated in a blocking reagent (5% BSA) for 2 hours, followed by overnight incubation at 4°C in a 1:1000 dilution of rabbit-m6A primary antibody. Afterwards, the membrane was washed four times for 10 minutes each in 1X TBST, then incubated with a 1:1000 dilution of goat anti-rabbit secondary antibody for 2 hours at room temperature. The membrane was washed again four times for 10 minutes each in 1X TBST. Finally, the membrane was developed using ECL and imaged with a Uvitech imager.

### Protein Estimation and Immunoblot

Hippocampi were isolated from P21 C57BL/6 mice and lysed in RIPA buffer (25 mM Tris-HCl, pH 7.4, 150 mM NaCl, 1% NP-40, 0.5% sodium deoxycholate, 0.1% SDS) supplemented with protease (Sigma; 1 µl/ml) and phosphatase (Sigma; 1 µl/ml) inhibitors. The lysate was centrifuged at 12,000 x g for 10 minutes at 4°C to pellet the debris. The supernatant was collected and the protein concentration was determined using BCA assay. First, a series of protein standards with known concentrations was prepared by diluting a stock solution of bovine serum albumin (BSA). The BCA working reagent was prepared by mixing reagents A and B in a 50:1 ratio, as per the manufacturer’s instructions. Subsequently, 200 µL of the BCA working reagent was added to each well containing the standards and samples. The plate was then incubated at 37°C for 30 minutes to allow for the colourimetric reaction to develop. After incubation, the absorbance of each well was measured at 562 nm using a microplate reader. The protein concentrations of the samples were determined by comparing their absorbance values to the standard curve generated from the BSA standards.

The protein samples with 4X Laemmli buffer (250 mM Tris-HCl, pH 6.8, 8% SDS, 40% glycerol, 0.02% bromophenol blue, 20% β-mercaptoethanol) were heated at 95 ° C for 5 minutes. Equal amounts of protein were loaded onto SDS-PAGE gel and were run in running buffer (25 mM Tris, 192 mM glycine, 0.1% SDS). 5 µg protein was loaded for Tubulin β1, 30 µg of protein was loaded for FTO, and 30 µg of protein was loaded for METTL3. The proteins from the gel were transferred to a PVDF membrane using a transfer buffer (25 mM Tris, 192 mM glycine, 20% methanol) at 65 V for 90 minutes. After transfer, the membrane was blocked in 5% BSA (Bovine Serum Albumin) in TBST (20 mM Tris-HCl, pH 7.6, 150 mM NaCl, 0.1% Tween-20) for 1 hour at room temperature. The membrane was incubated with primary antibody against Tubulin β1 (Sigma; Anti-mouse; 1:5000), FTO (Abcam; Anti-mouse; 1:1000), METTL3 (Abcam; Anti-rabbit; 1:1000), diluted in blocking buffer, overnight at 4°C. The membrane was washed three times, for 10 minutes each, with TBST, then incubated with HRP-conjugated goat anti-mouse secondary antibody against Tubulin β1 (1:10000) and FTO (1:5000) and with HRP-conjugated goat anti-rabbit secondary antibody against METTL3 (1:5000), diluted in the blocking buffer for 2 hours at room temperature. The membrane was washed again for three times, for 10 minutes each, with TBST. For FTO overexpression and knockdown conformations, GAPDH was used as an internal control with a concentration of 1:10000 for primary (Sigma; Anti-Rabbit) and 1:10000 for secondary (HRP-conjugated goat anti-rabbit secondary antibody). Finally, the protein bands were detected using an enhanced chemiluminescence (ECL) substrate, and the signal was captured using an imaging system (Fig. S8-S13).

### qRT-PCR and cDNA preparation

P21 C57BL/6 male mice were sacrificed, and their hippocampi were harvested in 600 µl of ice-cold TRIzol reagent. The tissue was triturated with an 18 mm-gauge syringe to create a single-cell suspension. The mixture was centrifuged at 1000 x g for 5 minutes at 4°C to pellet debris. The supernatant was collected, and 300 µl of room-temperature chloroform was added and mixed thoroughly. After incubating on ice for 5 minutes, the mixture was centrifuged at 18,000 x g for 15 minutes at 4°C to separate the aqueous phase. The aqueous layer was carefully collected, and 150 µl of room-temperature isopropanol was added and mixed thoroughly. The mixture was incubated at −80°C overnight and then centrifuged at 18,000 x g for 3 hours at 4°C to precipitate the RNA. The supernatant was decanted, and the pellet was washed with 600 µl of 70% ethanol at room temperature and centrifuged at 18,000 x g for 1 hour at 4°C. The RNA pellet was air-dried in a laminar flow hood and dissolved in 20 µl of nuclease-free water warmed to 70°C. The RNA was treated with DNase to ensure purity. A cDNA library was prepared using the SuperScript III cDNA preparation kit. All qRT-PCR primers were made using the ENCODE database, and PCR was performed using SYBR Green.

**Table 1.**
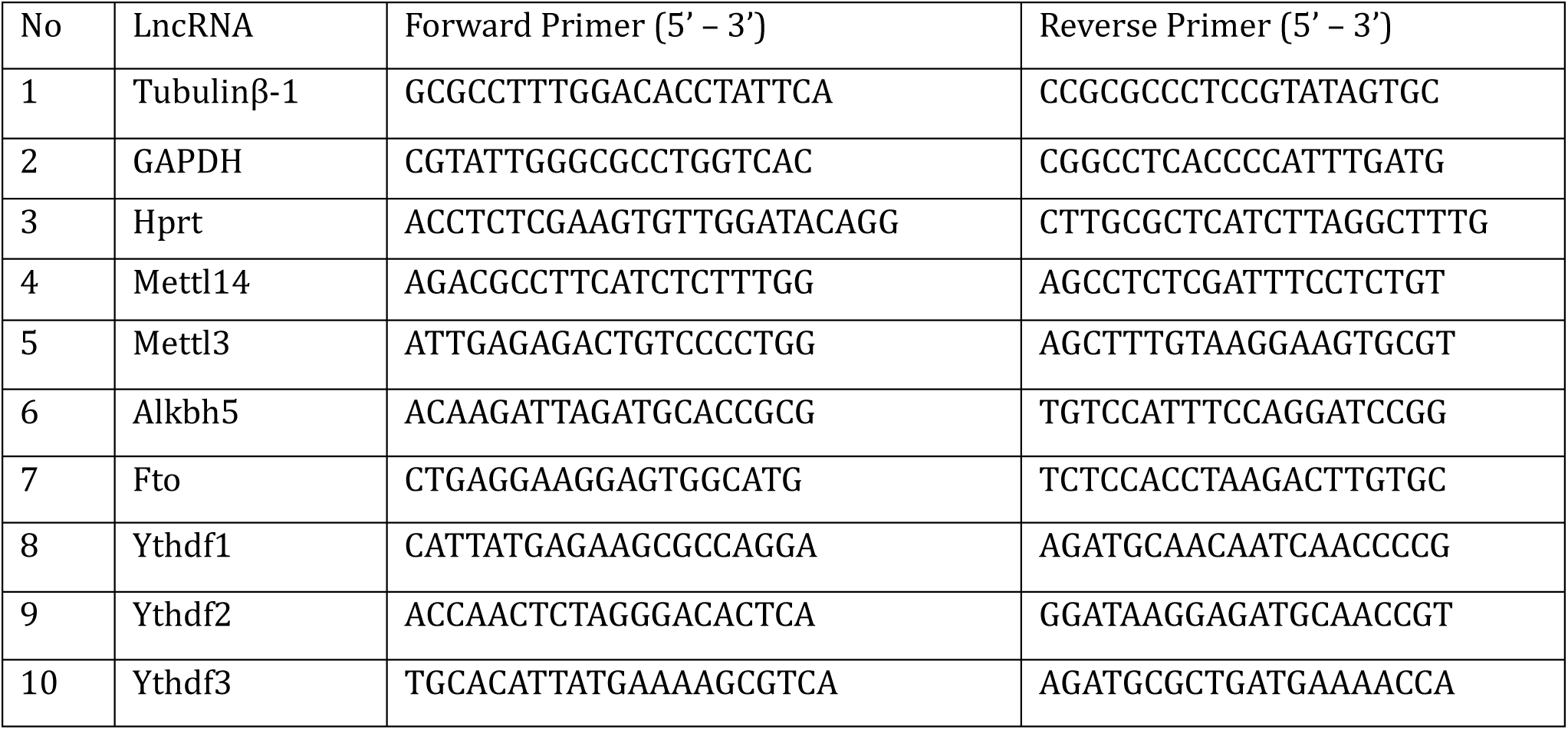
qPCR primers for m6A regulators.

### Cloning of FTO overexpression and knockdown vectors

FTO cDNA was obtained through PCR amplification from the pRK5-myc-FTO vector. The PCR product included an N-terminal T2A sequence and specific overhangs. The FTO-T2A PCR product was run on an agarose gel, after which the desired band was excised and purified. The purified FTO-T2A fragment was then cloned into the pAAV.hSynapsin.EGFP.WPRE.bGH (Osten-Frank) vector (Penn Core Vectors: p1696), which had been linearised using the AgeI restriction enzyme, through infusion cloning. To propagate the recombinant vector, the Stbl3 strain of E. coli was used (Fig. S2, S12).

For FTO RNAi, two independent shRNA constructs targeting FTO (shRNA1 and shRNA2) were individually cloned into the AAV-U6-sgRNA-hSyn-mCherry vector (Addgene plasmid #87916) using SacI and ApaI restriction enzyme sites. The use of two distinct shRNAs minimised the likelihood of off-target effects. A scrambled shRNA control was similarly cloned into the same vector to serve as a negative control (Fig. S6, S13).

**Table 2.**
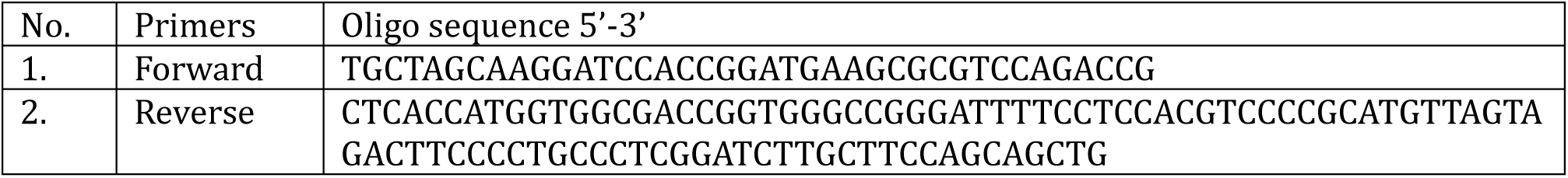
Infusion primers for insertion of FTO-t2a sequence in AAV-Synapsin-EGFP backbone.

**Table 3.**
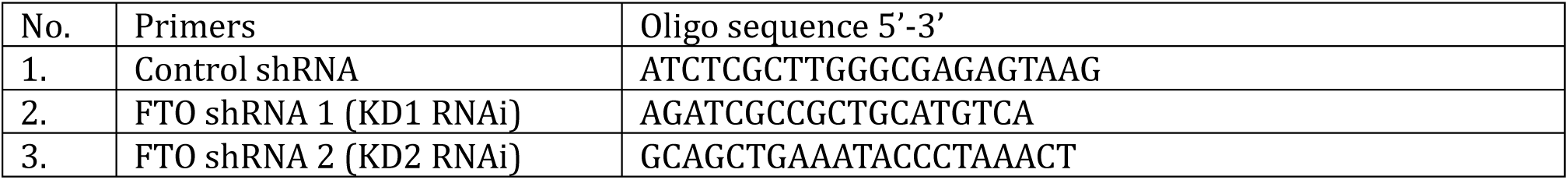
FTO shRNA primers inserted in AAV-U6-sgRNA-hSyn-mCherry backbone.

### HEK-293t cell culture and adeno-associated virus preparation

HEK-293t cells were grown and propagated in low-glucose DMEM solution (Gibco) with 10% fetal bovine serum. Cells were maintained in incubators at 37 °C with 5% CO2. All expression vectors were propagated utilising the Stbl3 strain of E. coli, while the packaging plasmids, pAAV2/9n and pDeltaF6, were propagated using the DH5α strain. Purification of the plasmid was achieved through an Endotoxin-free Maxiprep kit (Qiagen). HEK293t cells underwent transfection using the purified plasmid. For transfection of 2 × 10^6 cells (one 100mm plate), a mixture comprising 10 µg of the chosen expression vector, 10µg pAAV2/9n, and 20µg of pAdDeltaF6 was prepared in a microcentrifuge tube. To this mixture, 500 µl of 2X HBS was added, and the total volume was adjusted to 1ml with nuclease-free water. 50 µl of 2.5 M calcium chloride (CaCl2) was added dropwise, followed by vortexing and incubation at room temperature for 25 minutes. Post-incubation, the mixture was added dropwise to the culture in the 100mm plate and thoroughly mixed. The culture was then maintained at 37°C with 5% CO2 in low-glucose DMEM (Gibco) supplemented with 10% fetal bovine serum, with a single media change after 8 hours. The culture was maintained for 96 hours post-media change. The addition of NaCl (final concentration 500 mM) to the plate was followed by incubation at 37°C for 3 hours to lyse the cells and release the virus into the supernatant. The supernatant was collected in a centrifuge tube and stored at 4°C. The remaining cells in the dish were scraped and collected, undergoing a free-thaw cycle: −80°C for 30 minutes followed by 37°C for 30 minutes, repeated three times to ensure complete dissociation of viral particles from the cell membrane. The scrapped cells and the previously collected supernatant were filtered through a 100 µ nylon mesh filter initially to separate the membrane debris. The resulting filtrate was further filtered through a 45 µ filter, and the final filtrate was concentrated using Amicron-Ultra-15 (Merk Millipore #UFC910024) filters. Virus titre determination and calculation of the multiplicity of infection (MOI) were carried out by infecting 80K HEK293T cells with serially diluted AAV solutions. An MOI of 1-2 was employed for subsequent experiments.

### Primary Neuronal Culture

Following a previous protocol (Kaech & Banker, 2006), primary hippocampal culture was performed. Pregnant CD1 dams were selected and sacrificed in a CO2 chamber when their embryos were at the E16 stage (embryonic day 16). The E16 pups were dissected out from the uterus and kept in ice-cold Dissection Media for proper anaesthesia. The pups were then decapitated to extract the brain. The hippocampi were dissected and collected in cold dissection media (Gibco). It was then digested with the addition of 2.5% trypsin to the dissection media at 37 °C for 10 mins. Trypsin-treated tissue was washed with dissection media thrice at room temperature and collected in Glial Minimum Essential Media (GMEM; Gibco). Further, trituration was done first with an unpolished Pasteur pipette and then with a fire-polished Pasteur pipette to obtain a single-cell suspension, which was plated on poly-L-lysine-coated 60mm dishes (1 mg/mL, Sigma) at a concentration of 200-250 cells/mm^2^. The culture was maintained at 37 °C, 5% CO2 in Neurobasal Media (Gibco) with B27 supplement (Gibco) throughout the experiment. Cells were infected with AAV at DIV-7 and harvested at DIV-21 for further experiments.

### Stereotaxic Surgery

All surgical procedures were conducted on male P30 C57BL/6 animals. Anaesthesia was induced by a combination dose of Ketamine (80mg per 1kg of body weight) and Xylazine (20mg per 1kg of body weight), administered via subcutaneous injections. Following anaesthesia, the fur covering the head was shaved, and the animal was positioned onto the stereotaxic stage. An incision was made on the skin, exposing the underlying skull. The coordinates of the Bregma were precisely determined as (X, Y, Z) = (0, 0, 0), serving as a reference point. The skull was drilled to create a small aperture in the cranium at the predetermined coordinates for CA1 (Anterior −0.18cm; Lateral +/−0.19cm; Ventral −0.12cm relative to Bregma (0, 0, 0)). The viruses were bilaterally injected into the CA1 region at a controlled rate of 0.1 µl/min, with a total volume of 0.5 µl per hemisphere. Following the injection of 0.25 µl, a brief 5-minute pause was given to prevent excessive cranial pressure buildup. The syringe was carefully withdrawn 10 minutes after the 0.5 µl injection and an interval of 5 minutes was maintained before injection into the second hemisphere. Upon completion, the surgical site was sealed, and the incision was sutured using a nylon thread. The animal was transferred to a clean cage, where it was allowed to recover for 1 week. Experimental procedures were conducted 2 weeks post-surgery, ensuring ample recovery time.

### Cryosectioning

For cryosectioning, the mouse brain was harvested and fixed using a 4% paraformaldehyde solution to preserve tissue integrity. The brain was further incubated in a 4% paraformaldehyde solution for complete fixation of tissue. Post-fixation, the brain is incubated in a 30% sucrose solution until it becomes fully saturated, which takes 5-6 days. The brain is then flash-frozen using dry ice to ensure rapid and uniform freezing. Using the cryostat, the frozen brain is sectioned into 50 µm thick slices, at a maintained temperature of around −20°C. These sections are collected onto slides pre-coated with gelatine to enhance tissue adherence. The slides are stored at −80°C or processed immediately for further staining and microscopic examination.

### Spatial Object Recognition Assay

Male C57BL/6 mice at postnatal day 50 (P50) were housed individually under standard housing conditions. The mice underwent a one-week habituation period where they were gently handled for 2 minutes daily. The spatial object recognition assay involved five sessions, each lasting 10 minutes (Leger et al., 2013; Wimmer et al., 2012). The first four sessions were training sessions conducted on the same day, followed by test session 3 hours or 24 hours later. In Session 1, mice were introduced to an empty arena (12 cm x 15 cm) with spatial cues such as triangular and circular markings on one wall and alternating black and white stripes on another wall. This allowed the mice to explore and habituate for 10 minutes. During Sessions 2-4, two objects were placed at different locations in the arena, with a 2 cm gap between each object and the arena wall, allowing the mice to explore freely for 10 minutes per session. A 5-minute interval between sessions was used to clean the arena with 70% ethanol to remove any odours. The training involved pairs of mice in two separate arenas simultaneously. For the test session, conducted 3 hours or 24 hours after the training, one object (a cylindrical glass object) in one arena was moved to a new location, while no changes were made in the other arena, which served as a control. The mice were then reintroduced to their respective arenas and allowed to explore for 10 minutes. All sessions were recorded using an overhead-mounted Logitech HD webcam C270. The recordings were analysed using MATLAB-based software to measure the time spent with each object. Movement patterns and bout counts were determined, with a bout defined as an exploration of an object within a 2 cm radius, excluding climbing on the object. Additionally, all analyses were manually blind-tested.

The discrimination index (DI) was calculated to assess the animals’ ability to recognise object displacement, using the formula:

DI = [time spent with object displaced - time spent with object not displaced] / [time spent with displaced object + time spent with undisplaced object]

A positive DI indicated a preference for the displaced object, reflecting the retention of spatial memory.

### Light/Dark Box Assay

Male C57BL/6 mice at P42 underwent a one-week habituation period in a low-lux environment (2-5 Lux) (Bourin & Hascoe t, 2003; Takao & Miyakawa, 2006). Each mouse was gently habituated for two minutes daily in the experimental room during this period. Throughout the study, the mice were individually housed in standard cages under standard housing conditions. The experimental setup included a light/dark apparatus consisting of a dark box (occupying one-third of the total exploration area) and a bright box (occupying two-thirds of the total exploration area), separated by a sliding door. The dark box was kept at a low light intensity of 2 Lux, while the bright box had a higher intensity of 390 Lux. At the beginning of each test session, the mouse was placed in the dark box. After 10 seconds, the door separating the two boxes was opened, allowing the mouse to explore both compartments freely for 10 minutes. To track and analyse the mouse’s behaviour, a Logitech HD webcam C270 was positioned overhead to record its movements. The recorded data was analysed using Toxtrac, a free online software, which provided measurements such as the time spent in the light box, distance travelled within the light box, transitions between compartments, etc. (Rodriguez et al., 2018).

### Puromycin incorporation assay

To assess global protein synthesis in primary hippocampal neurons, the Click-iT™ Plus OPP Protein Synthesis Assay Kit, Alexa Fluor™ 488 (Thermo Fisher Scientific), was employed following the manufacturer’s protocol. Primary hippocampal neuronal cultures were prepared and maintained under standard conditions. At DIV 7, neurons were transfected with 2µg FTO knockdown constructs (shRNA1 and shRNA2) and Control RNAi using Lipofectamine 2000 (Invitrogen). To halt transcriptional activity, Actinomycin D (5 µg/mL) was applied to the culture media for 30 minutes at 37°C before OPP treatment to ensure that any newly synthesized proteins result from translation of pre-existing mRNA pools. Following actinomycin treatment, neurons were incubated with O-propargyl-puromycin (OPP) at a final concentration of 20 µM for 30 minutes at 37°C. OPP is incorporated into nascent polypeptide chains, acting as a puromycin analogue to label newly synthesised proteins. After incubation, cells were gently washed with PBS and fixed with 4% paraformaldehyde for 15 minutes at room temperature. Cells were permeabilized using 0.25% Triton X-100 in PBS for 10 minutes, followed by Click-iT reaction according to the manufacturer’s instructions to conjugate Alexa Fluor™ 488 azide to incorporated OPP. Nuclei were counterstained and were mounted using Vectashield-DAPI.

### Imaging

Fluorescence imaging was performed using a point-scanning confocal microscope (Nikon AXR). The integrated OPP signal per cell was quantified using ImageJ. Images were acquired with a 60X oil immersion objective (NA = 1.4) at a resolution of 1024 × 1024 pixels, using a step size of 0.75 µm across 10–12 z-sections per image. Fast sequential acquisition mode was used with minimum spectral crosstalk. Fluorophores were excited with 405 nm (DAPI), 488 nm (OPP incorporation/GFP), 561 nm (mCherry), and 640 nm (MAP2/far-red) lasers. For higher-magnification imaging of dendrites, a 2X optical zoom was applied with a step size of 0.5 µm. Post-acquisition, images were processed using Fiji, and maximum intensity projections were generated. mCherry-positive neurons were masked using MAP2 staining, and OPP intensity was measured within selected dendritic regions of interest (ROIs) to assess *de novo* protein synthesis. The same approach was used to quantify GFP intensity within the soma.

### Analysis of RNA-IP data

Raw data were obtained from the independent datasets. Adapter trimming and quality filtering were performed using Cutadapt (v3.7), where low-quality bases (Phred score <20) and Illumina adapter sequences were removed. The resulting high-quality reads were aligned to the mouse reference genome (mm10) using HISAT2 (v2.2.1), and alignment statistics were compiled. Peaks of enriched methylation were identified using exomePeak (v2.16.0), which also detected differential methylation between experimental conditions. Statistically significant and replicate-consistent peaks were classified as hypermethylated (log2FC > 0) or hypomethylated (log2FC < 0), with further filtering based on a ≥2-fold change. Peaks were annotated as coding or noncoding based on genomic context. Volcano plots were generated to visualise p-values and fold changes, highlighting robust differentially methylated sites across conditions. Violin plots were generated to visualise the global shift of m6A deposition along the transcripts.

### Primary neuronal culture recordings

Primary hippocampal neurons were prepared and maintained under standard conditions. Neurons were transfected at DIV7 with 2 µg of FTO-targeting shRNA constructs (shRNA1) or control RNAi using Lipofectamine 2000 (Invitrogen). Recordings were performed at DIV18–DIV25.

Whole-cell patch-clamp recordings were performed using borosilicate glass electrodes (3–8 MΩ). Neurons were voltage-clamped at −70 mV. mEPSCs were recorded for 300 s per cell in extracellular solution consisting of (in mM: NaCl 119, KCl 5, CaCl₂ 2, MgCl₂ 2, glucose 30, HEPES 10), with pH adjusted to 7.4 and osmolarity of 310–320 mOsm, supplemented with tetrodotoxin (1 µM) and bicuculline (10 µM). The internal solution consisted of (in mM: cesium gluconate 100, EGTA 0.2, MgCl₂ 5, ATP 2, GTP 0.3, HEPES 40), with pH adjusted to 7.2 and osmolarity of 285–290 mOsm.

Cells with series resistance ≤30 MΩ and stable recordings were analysed. Cells with holding currents < −100 pA or unstable recordings were excluded.

### Statistical analysis

Statistical analyses were performed for all experiments, with data presented as mean ± SEM. The designation ‘N’ refers to the number of pregnant dams or litters used, while ‘n’ indicates the total number of pups obtained from these dams. For qRT-PCR data, immunoblots on m6A regulators, and puromycin incorporation assay, we employed unpaired Student’s t-tests with Welch’s correction for unequal variance. Paired Student’s t-tests were used for analysing FTO overexpression and knockdown efficiency data in HEK293t cells. Two-way ANOVA with Fisher’s LSD test and no correction was applied for analysing light/dark and SOR data. Data for litter effect was assessed using 2-way ANOVA and corrected with Bonferroni’s multiple comparison test. Post-hoc power analysis was conducted using GPower 3.1 for all experiments. Outliers were identified using the ROUT method (0.1% aggressive). Immunoblot band intensity analysis and image stitching were performed using ImageJ software. The volcano plot was obtained with FDR > 0.05 Gene Ontology plots were generated using RStudio and SRplot (Tang et al. 2023). Statistical significance was defined as p < 0.05.

## Results

### Maternal separation induces deposition of m^6^A mark in hippocampal transcriptome during early development

Previous studies established the role of m^6^A in developing neural cells and also identified its dysregulation as an emerging hallmark of neurodegenerative disorders, e.g. Huntington’s Disease (Du et al., 2019; Pupak et al., 2022). To understand the effect of ELS on global m^6^A modification, we performed a bulk RNA-seq from m^6^A-modified precipitated hippocampal RNAs (Fig. 1A-B). This approach revealed 291 significantly hypermethylated transcripts among 853 hypermethylated candidates (both mRNAs and lncRNAs) and one significantly hypomethylated transcript out of 71 hypomethylated candidates (Fig. 1C). Moreover, analysis of the distribution of methylation peaks indicated a broader shift in m^6^A deposition along lncRNA transcripts and within the 3’-UTR and 5’-UTR of mRNAs, suggesting a mechanism by which early-life stress may reprogram hippocampal gene expression through RNA methylation (Fig. 1D-E). In addition, we examined the m^6^A peak enrichment around canonical DRACH motifs within transcripts encoding key RNA-binding proteins (RBPs) relevant to neuronal function. Among the candidates analysed exhibited reduced enrichment at the adenosine of the DRACH motif, where the m^6^A peak is typically localised (Fig. 1H). A comparable pattern of altered distribution was observed for Pum1 and, to a lesser extent, G3bp2 (Fig 1M, 1I). In contrast, Srsf2, Srsf9 and Fxr1 did not display detectable changes in peak profiles (Fig. 1L, 1K, 1J). Whereas Srsf1 and Pum2 showed a shift in the m6A peak (Fig. 1F, 1G), thereby suggesting the existence of distinct modes of stress-induced remodelling of the RBP-associated methylation sites.

**Figure 1.**
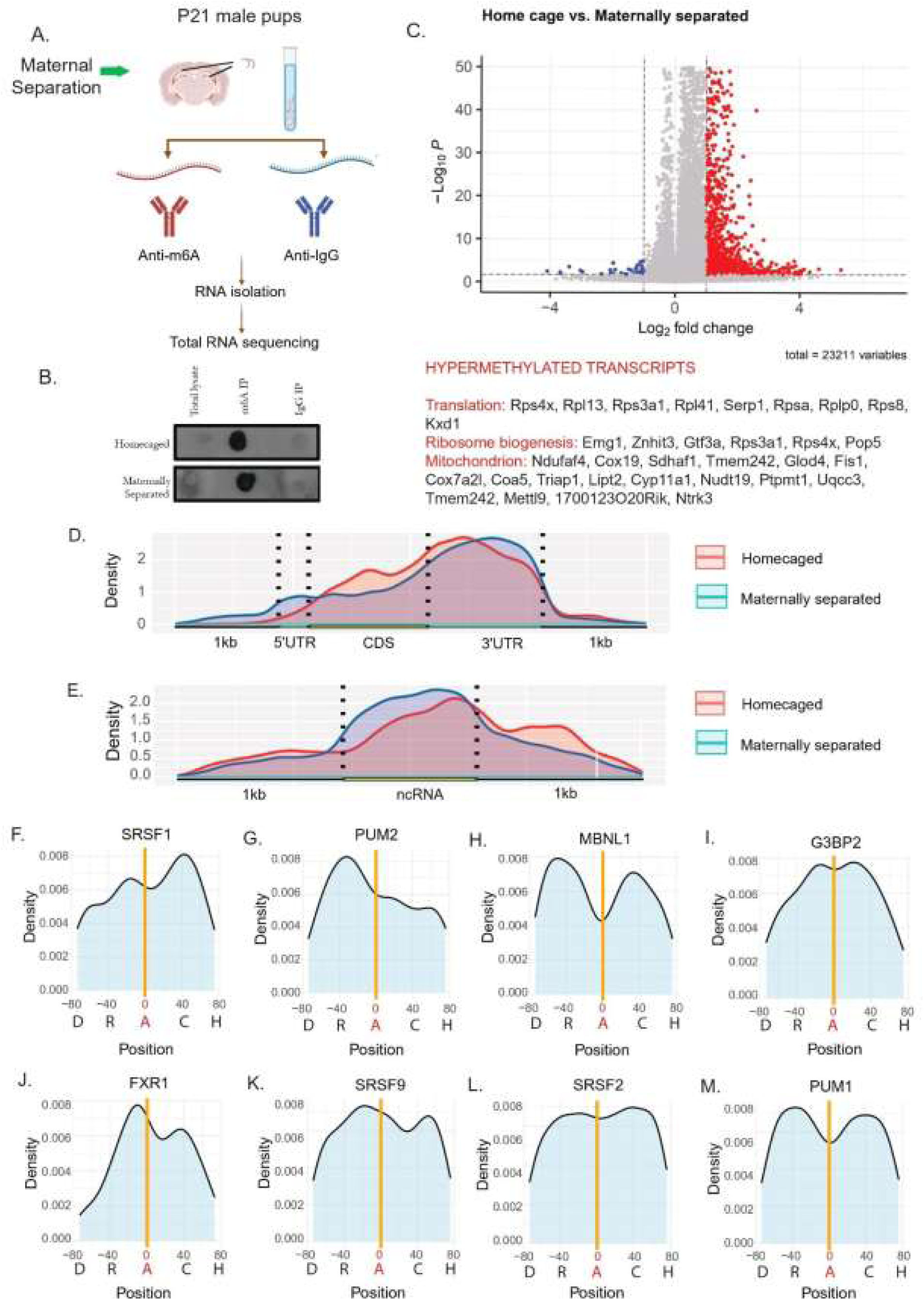
Maternal Separation enhances m^6^A modifications of hippocampal transcripts. **(A)** Experimental workflow, N = 3. **(B)** Representative dot blot showing m6A immunoprecipitation (IP) efficiency, N = 3. **(C)** Volcano plot for RNA expression of home-caged (HC) versus maternally separated (MS) P21 pups. Red dots = significantly upregulated RNAs; blue dots = significantly downregulated RNAs in maternally separated pups, total variables = 23,211, N = 3, FDR > 0.05. **(D)** Density plot for distribution of m^6^A peaks along different regions of mRNA transcripts (5’ UTR, coding sequence (CDS), and 3’ UTR), between HC and MS groups, N=3. **(E)** Density plot for distribution of m^6^A peaks along lncRNAs in HC vs MS groups, N=3. **(F-M)** The relative change position of the binding site of different RBPs: **(F)** SRSF1, **(G)** PUM2, **(H)** MBNL1, **(I)** G3BP2, **(J)** FRX1, **(K)** SRSF9, **(L)** SRSF2, and **(M)** PUM1 in relation to the m^6^A (DRACH) motif; the “A” position is considered as 0. *Also see Supplementary Figure 1*.

### Maternal separation selectively downregulates FTO expression in a sex-specific manner

Given the dynamic regulation of m^6^A by readers, writers and erasers, we examined their expression at the transcript as well as protein level, to identify the factors underlying the altered methylation landscape induced by maternal separation.

We assessed the expression of core m^6^A regulatory components in hippocampal tissue from P21 male mice using quantitative real-time PCR. No significant alterations were detected in the methylases METTL3 (52.93±41.65, p=0.24) and METTL14 (65.63±51.07, p=0.24) (Fig. 2A-B), the demethylase Alkb5 (29.07±18.19, p=0.15) (Fig. 2D), or the readers YTHDF2 (14.91 ± 19.46, p=0.46) and YTHDF3 (17.57± 14.15, p=0.24) (Fig. 2F-G). In contrast, maternal separation significantly reduced the expression of the demethylase FTO (38.58 ± 14.64, p=0.02) (Fig. 2C) and the reader YTHDF1(36.15 ± 14.35, p=0.03) (Fig. 2E), suggesting that early-life stress selectively targets components of the m^6^A machinery.

**Figure 2.**
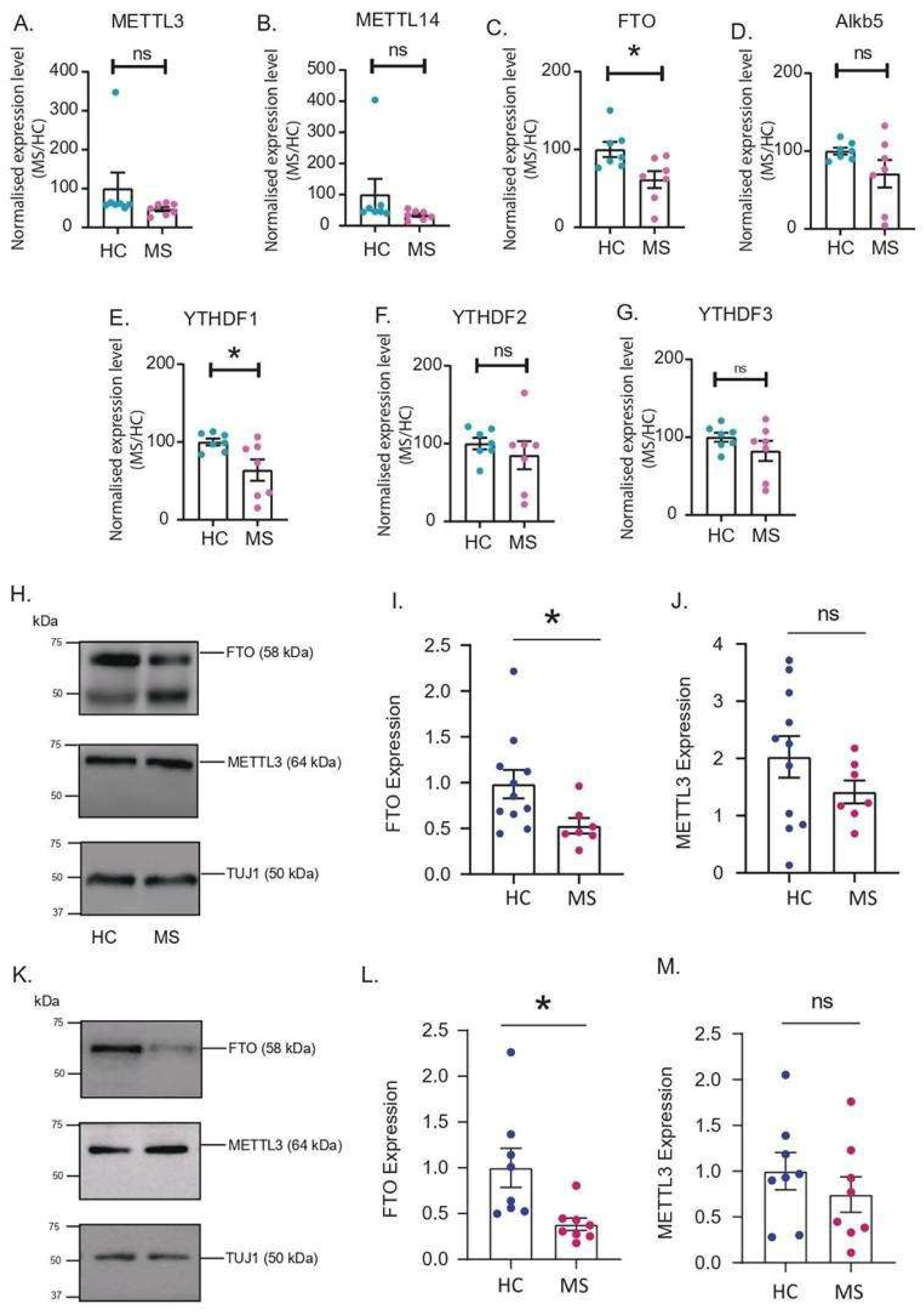
Regulation of readers, writers and erasers upon maternal separation. Differential expression of methylases **(A)** METTL3 and **(B)** METTL14, demethylases **(C)** FTO and **(D)** Alkb5 and reader proteins **(E)** YTHDF1, **(F)** YTHDF2 and **(G)** YTHDF3 transcripts as obtained by qPCR. *p < 0.05, **p < 0.01, ns = not significant. N=7, Data shown as Mean ± SEM. Unpaired Student’s t-test with Welch’s correction. Immunoblot showing expression of FTO and METTL3 in males **(H)** at P21, and **(K)** at P28. Quantification of FTO expression in **(I)** P21; N=3-4, n=7-11 and in **(L)** P28; N=3-4, n=8, males. Quantification of METTL3 expression in **(J)** P21 and in **(M)** P28 males. *p < 0.05, **p < 0.01, ns = not significant. Mean ± SEM. Unpaired Student’s t-test with Welch’s correction.

To observe the expression at the protein level, we performed immunoblotting for METTL3 and FTO in hippocampus lysates. In P21 male mice, METTL3 protein abundance was not significantly altered (0.6140 ± 0.4141, p=0.15) (Fig. 2J), whereas FTO expression was significantly reduced following MS (0.4549 ± 0.1757, p=0.02) (Fig. 2I). To determine whether this effect persisted into young adulthood, we re-examined the expression profile of METTL3 and FTO at P28. We found a similar pattern where FTO was significantly downregulated (0.6170±0.2238, p=0.023) and there was no effect on METTL3 protein expression (0.2554±0.2805, p=0.37) (Fig 2L, M).

To understand the sexual dimorphism in this regulation, we also examined the levels of FTO and METTL3 in the hippocampus of maternally separated female mice at P21 and P28. At P21, expression of FTO as well as METTL3 in female hippocampus were similar to that of P21 males; with a significantly downregulated FTO (0.6842±0.178, p=0.004) and insignificant change in METTL3 expression (0.098±0.188, p=0.612) (Fig. S1A-C). But at P28, there was no significant difference in expression of FTO (0.049±0.316, p=0.87) or METTL3 (0.0005±0.33, p=0.99) between the MS and the control mice (Fig. S1D-F). The sustained reduction of FTO, observed in males, appears to be transient in females. This highlights a sexually dimorphic regulation in epigenetic response to ELS.

### FTO overexpression does not rescue MS-induced anxiety-like behaviour

MS markedly increased anxiety in P45 C57BL/6 male mice, as shown by reduced distanced traveled in the light compartment (5.908±2.796, p=0.04) and fewer transitions made between the two compartments (13.86±3.683, p=0.0009) as compared to the HC (homecaged) animals (Fig. 3A-3D). The change in time spent in the light compartment showed no significant difference (19±72.26, p=0.79) between the MS and the HC cohort. This was probably because, due to heightened anxiety, the MS animals froze upon entering the light box (Fig. 3E).

**Figure 3.**
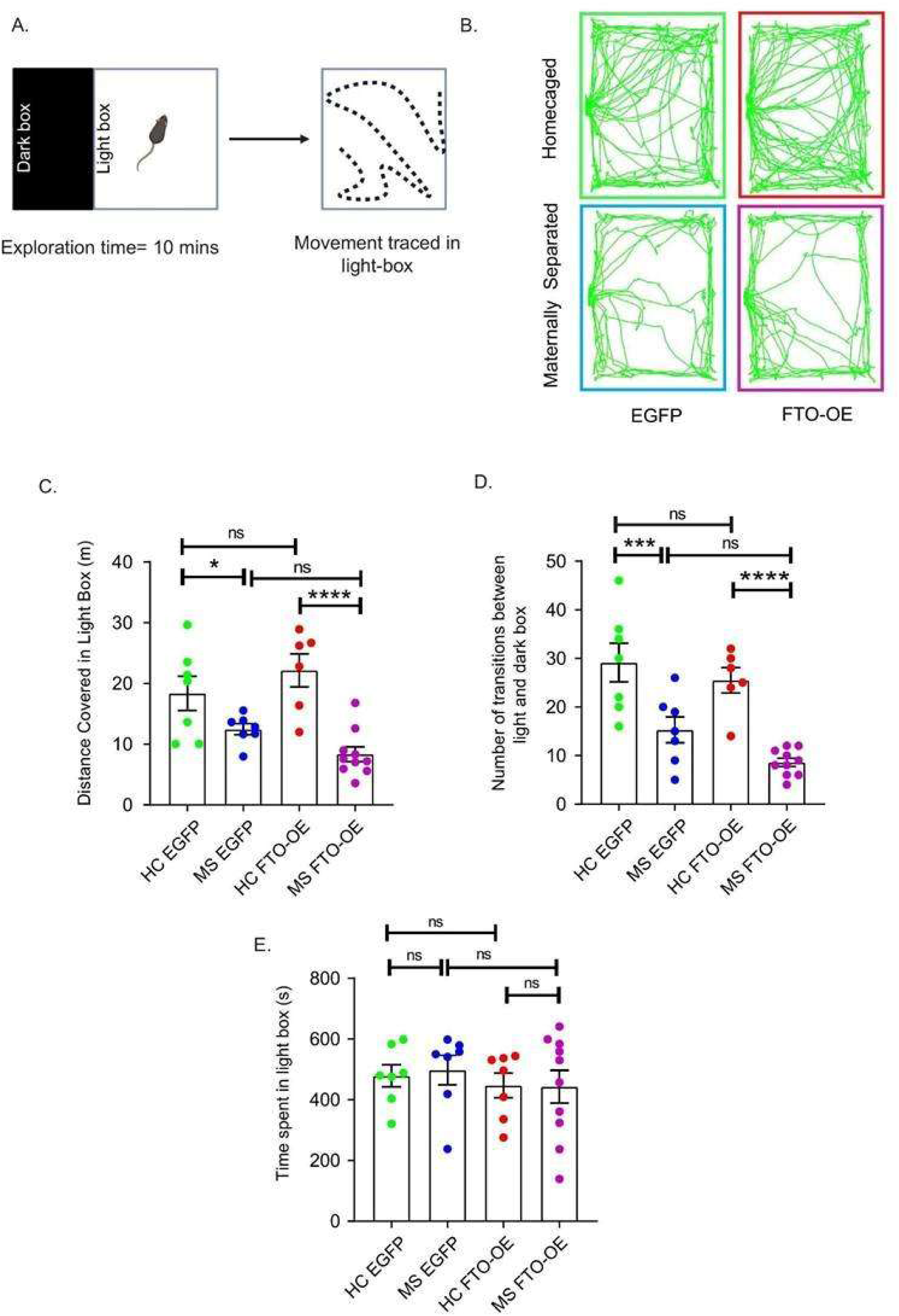
FTO overexpression does not reverse elevated anxiety following maternal separation. **(A)** Paradigm depicting light/dark box assay. **(B)** Representative trace in the bright-lit arena in the light/dark box. HC = Homecaged, MS = Maternally Separated, EGFP = Empty plasmid with GFP fluorescence, FTO-OE= Plasmid with FTO overexpression and GFP fluorescence. **(C)** Distance covered in the light box (in meters). **(D)** Total number of transitions made between light and dark box. **(E)** Time spent in the light box (in seconds). Data shown as mean±SEM, N=3, n=7-10, *p<0.05 **p< 0.01, ***p<0.005, ****p<0.0001, ns not significant. Two-way ANOVA with Fisher’s LSD test. *Also see supplementary figure S5*

FTO overexpression in the hippocampus did not alleviate anxiety-like behaviour in the MS group. FTO-OE (FTO overexpression) MS mice exhibited comparable anxiety levels to EGFP MS controls, with no significant differences in time spent in the light box (54.47±66.62, p=0.42), distance travelled (4.107±2.578, p=0.12), or transitions made (6.686±3.395, p=0.059). FTO overexpression alone did not affect baseline anxiety, as no differences were observed between FTO-OE HC and EGFP HC mice across time (46.57±75.21, p=0.54), distance (3.796±2.91, p=0.2) or transitions (3.643±3.833, p=0.35) (Fig. 3C-E). FTO-OE MS mice displayed elevated anxiety relative to FTO-OE HC controls (time: 11.1±69.81, p=0.87; distance: 13.81±2.701, p< 0.0001; transitions: 16.9±3.558, p< 0.0001), which again mirrored the behavioural pattern of EGFP MS mice (time: 11.1±69.81, p=0.87; distance: 13.81±2.701, p< 0.0001; transitions: 10.21±3.833, p=0.01) (Fig. 3C-E). To rule out litter-specific bias, we analysed behavioural variations within litters under the same conditions and detected no significant differences (Fig. S5). Collectively, FTO overexpression in the hippocampus did not rescue the anxiety phenotype induced by MS.

### FTO overexpression rescues spatial memory deficits of young adult mice caused by maternal separation

FTO regulates memory consolidation via an m^6^A-dependent BDNF-TrkB signalling pathway (Chang et al., 2023; Song et al., 2024). Previous studies have demonstrated that de novo protein translation is differentially engaged across early (1–3 hrs) and late (24 hrs) phases following neuronal activity, corresponding to short-term and long-term memory formation (Santini et al., 2014, Schafe et al., 2000). We therefore investigated the role of FTO in rescuing spatial memory deficits induced by ELS by testing spatial memory at two time points post-training: 3 hours and 24 hours.

Test at 3 hours post-training: The SOR assay measures preference for a displaced object over a non-displaced object. But at 3 hours post-training, the percentage time spent with displaced object in test session of EGFP-MS animals was comparable to that of the EGFP-HC group (3.739±6.03, p>0.0009). Additionally, MS mice receiving the FTO-OE (FTO-OE MS) construct also did not display a significant difference in percentage time spent with displaced object in test session compared to their EGFP-MS counterparts (5.028±6.03, p>0.0009) (Fig. 4C). Briefly, no significant change in spatial memory was observed in MS animals upon FTO overexpression at the hippocampus 3 hours post training.

**Figure 4.**
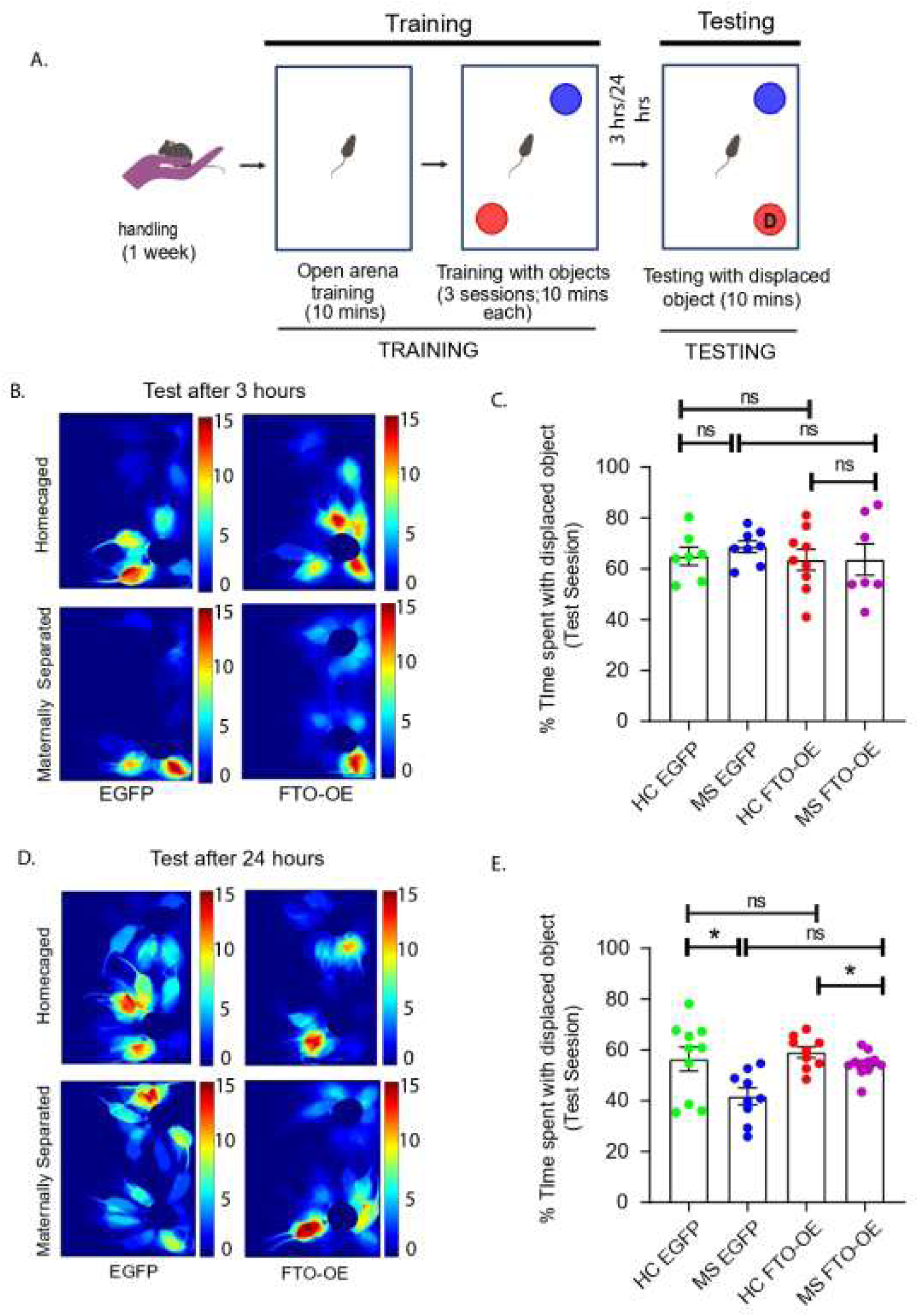
FTO overexpression reverses the effect of MS-induced deficits in memory. **(A)** Paradigm for SOR. **(B, D)** Representative heat maps in the arena with the displaced object during the test session, white ‘D’= object displaced; **(B)** 3 hours and **(D) 24** hours post-training. HC = Homecaged, MS = Maternally Separated, EGFP = Empty plasmid with GFP fluorescence, FTO-OE = Plasmid with FTO overexpression and GFP fluorescence**. (C, E)** Percentage time spent with displaced object during the test session, **(C)** 3 hours, N=3, n=7-9, and **(E)** 24 hours, N=3, n=9-11, post-training. Data shows Mean ± SEM., *p< 0.05, ns = not significant. Two-way ANOVA with Bonferroni’s correction. *Also see supplementary figures S3 and S4*.

Test at 24 hours post-training: The test sessions conducted 24 hours post-training showed significantly low exploration of the displaced object in EGFP-MS animals as compared to the EGFP-HC animals (14.67±4.508, p=0.051). When FTO was overexpressed in the hippocampus of MS animals (FTO-OE MS), this loss of spatial memory was rescued, as shown by a higher percentage of time spent with the displaced object in the test session as compared to EGFP-MS group (12.64±4.41, p=0.04). The exploration of FTO-OE MS animals was comparable to FTO-OE HC animals (4.692±4.41, p>0.0009) and to EGFP-HC animals (2.03±4.287, p>0.0009). Thus, FTO overexpression conferred a measurable improvement in spatial memory under ELS. Importantly, FTO-OE HC group did not alter baseline exploration with percentage time spent with displaced object in test session values similar to EGFP-HC mice (2.662±4.508, p>0.0009) (Fig. 4E).

The behavioural variation within litter was ruled out by studying the litter effect within the same experimental condition (Fig. S4). Further, no object preference was seen in any experimental group, ruling out any object bias (Fig. S3). Therefore, we observed that FTO overexpression in the hippocampus of MS-induced animals was sufficient to rescue spatial memory deficits 24 hours post-training.

### FTO downregulation reduces protein synthesis in primary hippocampal neurons

FTO knockdown was used to mimic the stress-induced reduction observed *in vivo*. This resulted in a significant decrease in puromycin-labelled nascent proteins, reflecting diminished translation activity in the absence of FTO (Fig. 5A). To minimise off-target effects, two shRNA constructs were used: FTO-RNAi_1 and FTO_RNAi_2. The knockdown efficiency of these constructs was verified by transfecting each construct and a control RNAi in HEK293T cells, followed by an immunoblot (FTO-RNAi_1: 55.12±20.87, p=0.02; FTO-RNAi_2: 64.84±22.78, p=0.15) (Fig. S6). FTO knockdown resulted in lower protein synthesis both in soma (FTO-RNAi_1: 54.89 ± 12.50, p<0.0001; FTO-RNAi_2: 50.17 ± 13.08, p=0.0003) and the dendrites (FTO-RNAi_1: 38.65 ± 7.579, p<0.0001; FTO-RNAi_2: 42.77 ± 7.259, p<0.0001) (Fig. 5B, C). Together, these data indicate that FTO is required to maintain basal protein synthesis in hippocampal neurons, and its loss disrupts translation in both somatic and dendritic domains of mature neurons.

**Figure 5.**
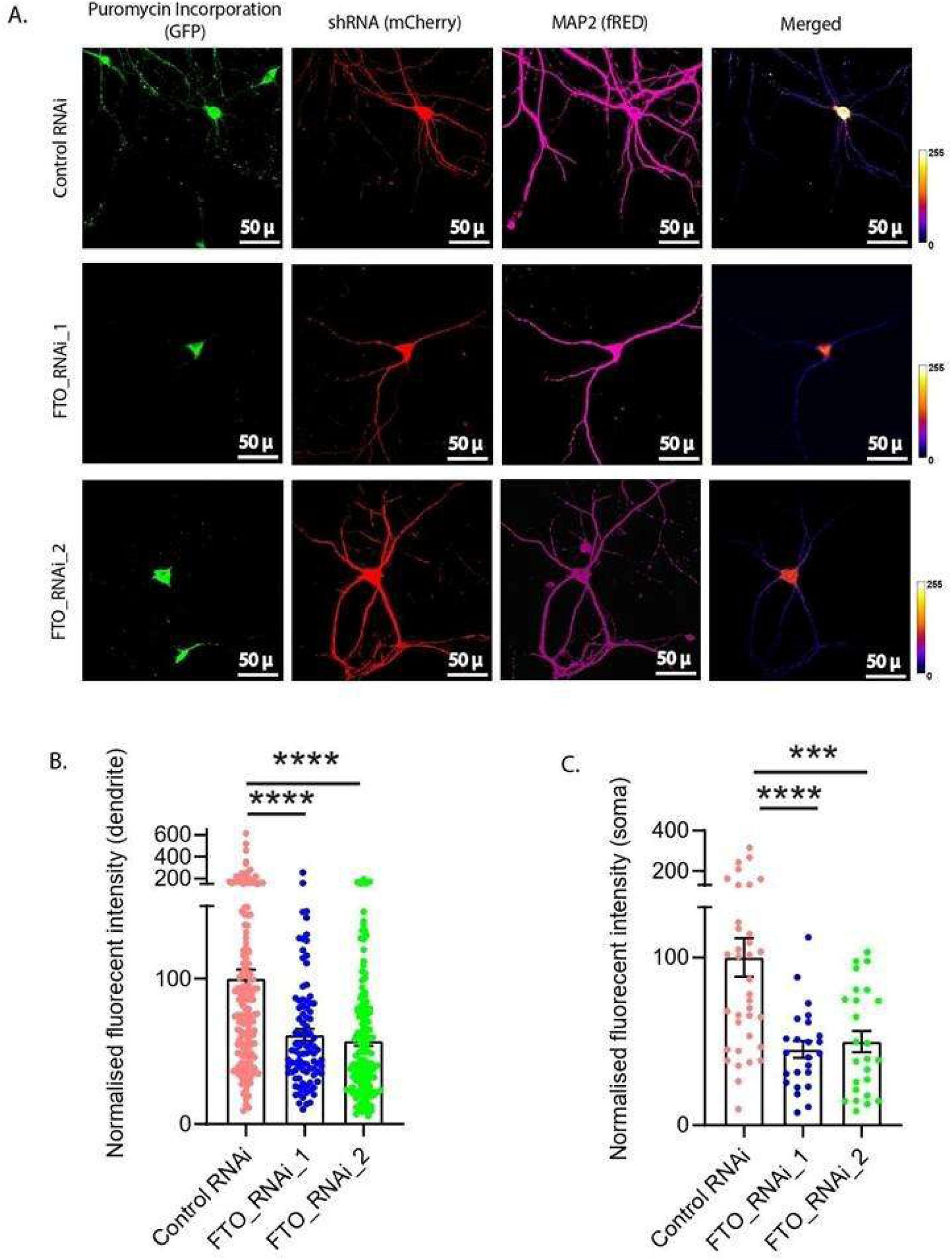
FTO RNAi reduces translation in primary hippocampal neurons. **(A)** Representative images of cultured primary hippocampal neurons transduced with control or FTO-targeting shRNA constructs (FTO_RNAi_1 and FTO_RNAi_2), co-expressing mCherry (shRNA marker, red), O-propargyl-puromycin (OPP) (GFP channel, green), and MAP2 (fRed, magenta). Scale bars: 50 μ**. (B–C)** Quantification of puromycin incorporation as % fold change in signal intensity in **(B)** dendrites and **(C)** soma. Data are shown as Mean ± SEM. Statistical significance was determined by Unpaired t-test with Welch’s correction, N = 24-36, n = 94-172, ***p < 0.001, ****p < 0.0001.

### m^6^A-modified RNA -interacting RBPs regulate RNA metabolism including protein synthesis

Gene Ontology (GO) enrichment of the hypermethylated transcripts revealed broad associations with RNA metabolic processes, including RNA 5’-end processing, ribosome biogenesis, translation and ribonucleoprotein complex assembly (Fig. 6). To gain deeper insights into the functional pathways potentially regulated by these transcripts, we performed GO analysis on their interacting RBPs. The biological process showed a significant regulation for pathways including mRNA metabolic processes, RNA localisation, cytoplasmic translation and post-transcriptional regulation of gene expression. Other terms, like dendritic transport of ribonucleoprotein, highlighted the potential regulation of translation in neuronal processes. Additionally, enrichment of terms like stress granule assembly and 3’UTR-mediated RNA stabilisation and destabilisation suggest a possible MS-induced translational control of these RBPs (Fig. S7A).

**Figure 6.**
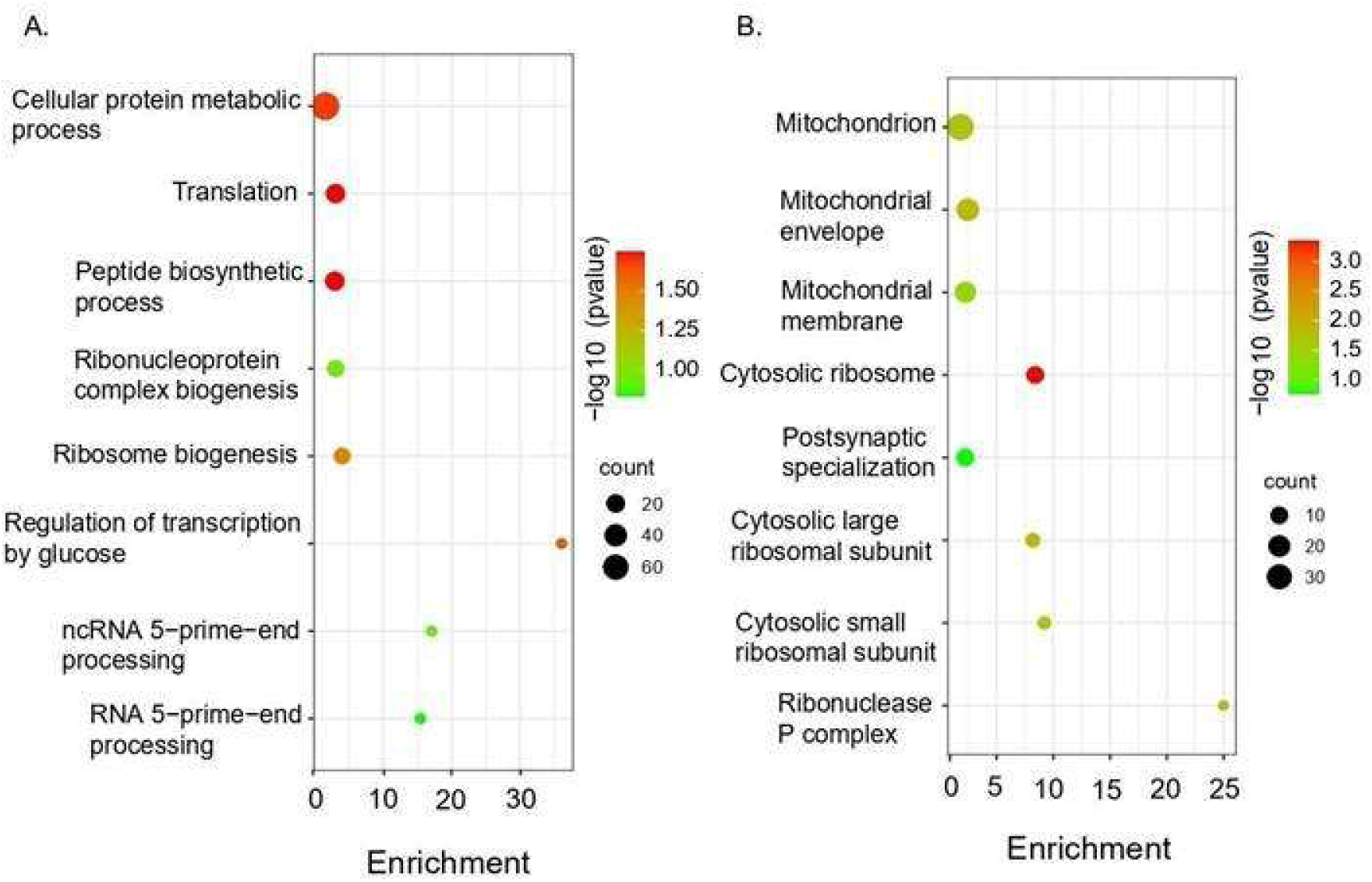
Gene Ontology enrichment of m⁶A-modified transcripts upon maternal separation. **(A)** Biological process, **(B)** Cellular components, FDR>0.05. *Also see supplementary figure S7*.

Further, cellular component analysis was enriched with terms including dendritic spines, distal axons, synapses, polysomes and cytoplasmic stress granules. These terms are associated with the localisation of these RBPs to neuronal compartments specialised for RNA storage and translation (Fig. S7B). Molecular function enrichment revealed a predominance of RNA-binding activities, including mRNA 3’UTR binding, translation initiation factor binding and N6-methyldenosine (m6A)-modified RNA binding, as well as interactions with regulatory RNA elements such as G-quadruplexes and miRNAs (Fig S7C). Together, these findings suggest that the m6A-modified RNA-interacting RBPs are well positioned to orchestrate spatial events at the synapse, potentially contributing to experience-dependent synaptic plasticity.

## DISCUSSION

The differential m^6^A methylation patterns observed in the hippocampus of male mice following maternal separation (MS) indicate a substantial reorganisation of the m^6^A epitranscriptomic landscape. Transcriptomic analysis of RNA-seq data revealed 291 transcripts that were significantly hypermethylated and only one that was hypomethylated after MS, suggesting a global shift toward hypermethylation (Fig. 1C). Previous studies have reported that m^6^A modifications within the 3′UTR promote transcript localisation to dendrites and axons in cultured hippocampal neurons (Livneh et al., 2020; Widagdo et. al., 2018). This process is essential for synaptic function and plasticity (Malovic & Pandey, 2023). Consistent with these observations, analysis showed a shift in m^6^A deposition toward the 3′UTR regions of coding transcripts and along the distal ends of lncRNAs, suggesting ELS-induced remodelling of the m6A-topology (Fig. 1D-E). We also observed a few candidates that are known to have a role in synaptic functions and translation, like Srsf1, Srsf2, Srsf9, Pum1, Pum2, Frx1, G3bp2 and Mbnl1. This shows a shift in m^6^A density along the conserved DRACH motif (D= A, G, U; R= A, G; H= A, C, U) that specifically recruits methyltransferase complexes to catalyse m6A modifications (Bayoumi & Munir, 2021; Ji et al., 2024; Santos-Rodriguez et al., 2024) (Fig. 1F-M).

Concurrently, the m^6^A demethylase FTO was significantly downregulated after MS, whereas the expression of methyltransferases remained unchanged in both sexes (Fig. 2, Fig. S1). The reduction in FTO levels following maternal separation aligns with findings from previous studies, which also report decreased FTO expression in the hippocampus in response to stress (Jiang et al., 2023; Kisliouk et al., 2020). However, FTO downregulation persisted longer in males than in females, hinting towards a sexually dimorphic regulation of m^6^A-decoration in the hippocampal post-ELS (Fig. 2, Fig. S1). The reduction of FTO may lead to the accumulation of m^6^A marks on stress-sensitive transcripts and potentially alter their stability, localisation, or translation. Thereby, it can impair neuronal and behavioural functions.

The reduction in FTO levels following MS is supported by previous findings, which report decreased FTO expression in the hippocampus and impaired working memory (Spychala & Ruther, 2019). Similarly, in individuals with major depressive disorder (MDD), FTO was found to be downregulated in the hippocampus (Mitsuhashi & Nagy, 2023). Recent studies have also shown that FTO expression at the synapse can affect coding as well as non-coding RNAs involved in synaptic functions and behaviour (Liau et al., 2023; Walters et al., 2017). Behaviourally, overexpression of FTO in the hippocampus did not reverse the anxiety-like phenotype induced by MS. The movement patterns were characterised by thigmotaxis and freezing, which are indicative of anxiety rather than exploration (Simon et al., 1994) (Fig. 3). These effects were not attributable to litter variability (Fig. S5). Thus, FTO overexpression appears insufficient to mitigate MS-induced anxiety, implying that additional stress-responsive molecular pathways contribute to anxiety-related behaviours. In contrast, FTO overexpression partially restored spatial memory deficits induced by MS in the SOR assay. While MS animals failed to recognise the displaced object at 24 hours post-training, FTO-overexpressing MS animals successfully identified it (Fig. 4D-E). The rescue effect was independent of litter differences (Fig. S4) and highlights that m^6^A-dependent regulation of memory-related transcripts is partially reversible by elevating FTO levels. Interestingly, there was no memory deficit in any condition 3 hours post-training (Fig. 4B-C). The absence of memory deficits 3 hours post-training indicates that short-term memory formation remains unaffected by MS and is independent of FTO expression. However, the impairment observed at 24 hours and its partial rescue by FTO overexpression imply that FTO influences the stabilisation of transcripts required for long-term memory persistence.

In primary hippocampal neurons, knockdown of FTO was done to mimic post-stress FTO levels in the hippocampus. We observed that loss of FTO led to a decrease in global protein synthesis both in dendrites and soma (Fig. 5). This implies that FTO modulates post-transcriptional translation efficiency rather than local translation. Therefore, FTO affects transcripts necessary for sustaining long-term synaptic plasticity. Stress-induced downregulation of FTO may impair memory consolidation by disrupting m^6^A-dependent regulation of protein synthesis in hippocampal neurons.

Further, to identify biological pathways affected by altered methylation, GO analysis of hypermethylated transcripts revealed enrichment in translation, ribosome biogenesis, and mitochondrial function, implying widespread effects on neuronal energetics and protein synthesis (Fig. 6). Further, GO analysis of RNA-binding proteins (RBPs) interacting with hypermethylated transcripts identified key terms related to RNA transport, localization, and stability, as well as dendritic localization, ncRNA processing, and 3′UTR-mediated RNA stabilization. Cellular component terms included synapse, dendrite, polysome, and P-body, while molecular function categories included translation activation, RNA and lncRNA binding, miRNA binding, and ribosome association (Fig. S7). These results collectively suggest that stress-induced hypermethylation perturbs RNA-RBP interactions critical for synaptic plasticity.

Together, these findings propose a model wherein early-life stress downregulates FTO in the hippocampus, leading to persistent hypermethylation of m^6^A-marked transcripts involved in protein synthesis and synaptic regulation. This epitranscriptomic shift alters RNA-RBP interactions and translation dynamics. Therefore, it can have long-lasting effects on neuronal plasticity and cognitive function. While anxiety-like behaviours appear independent of FTO-mediated demethylation, spatial memory deficits are, at least partially, reversible through restoration of FTO activity. This finding also highlights a dissociation between molecular pathways underlying stress-induced anxiety and memory impairments.

## Acknowledgements

The study is supported by NBRC core fund. Plasmids used for designing viral vectors were obtained from Addgene and Penn Vector Core. All art has been created with BioRender.com.

