## Supplemental Information for "FTO-dependent m6A RNA dysregulation underlies memory deficits induced by early-life stress"

**SUPPLEMENTARY DATA**

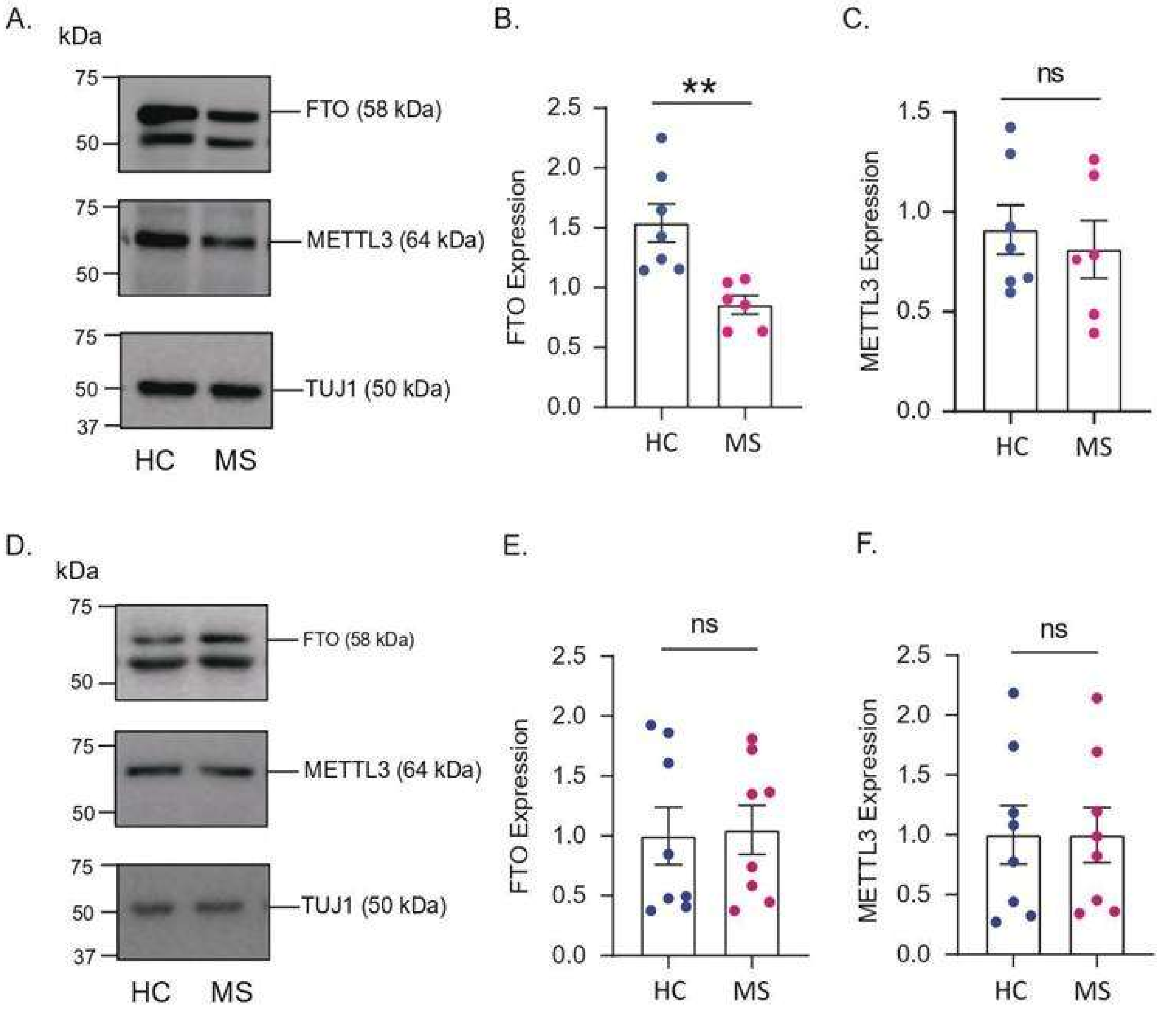


**Supplementary Figure 1. Expression of writers and erasers upon maternal separation in females.** Immunoblot showing expression of FTO, METTL3 and Tuj1 in females **(A)** at P21, and **(D)** at P28. Quantification of FTO expression in **(B)** P21; N=3-4, n=6-7 and in **(E)** P28; N=3-4, n=8, females. Quantification of METTL3 expression in **(C)** P21, N=3-4, n=6-7, and in **(F)** P28; N=3-4, n=8 males. *p < 0.05, **p < 0.01, ns = not significant. Mean ± SEM. Unpaired Student's t-test with Welch's correction.

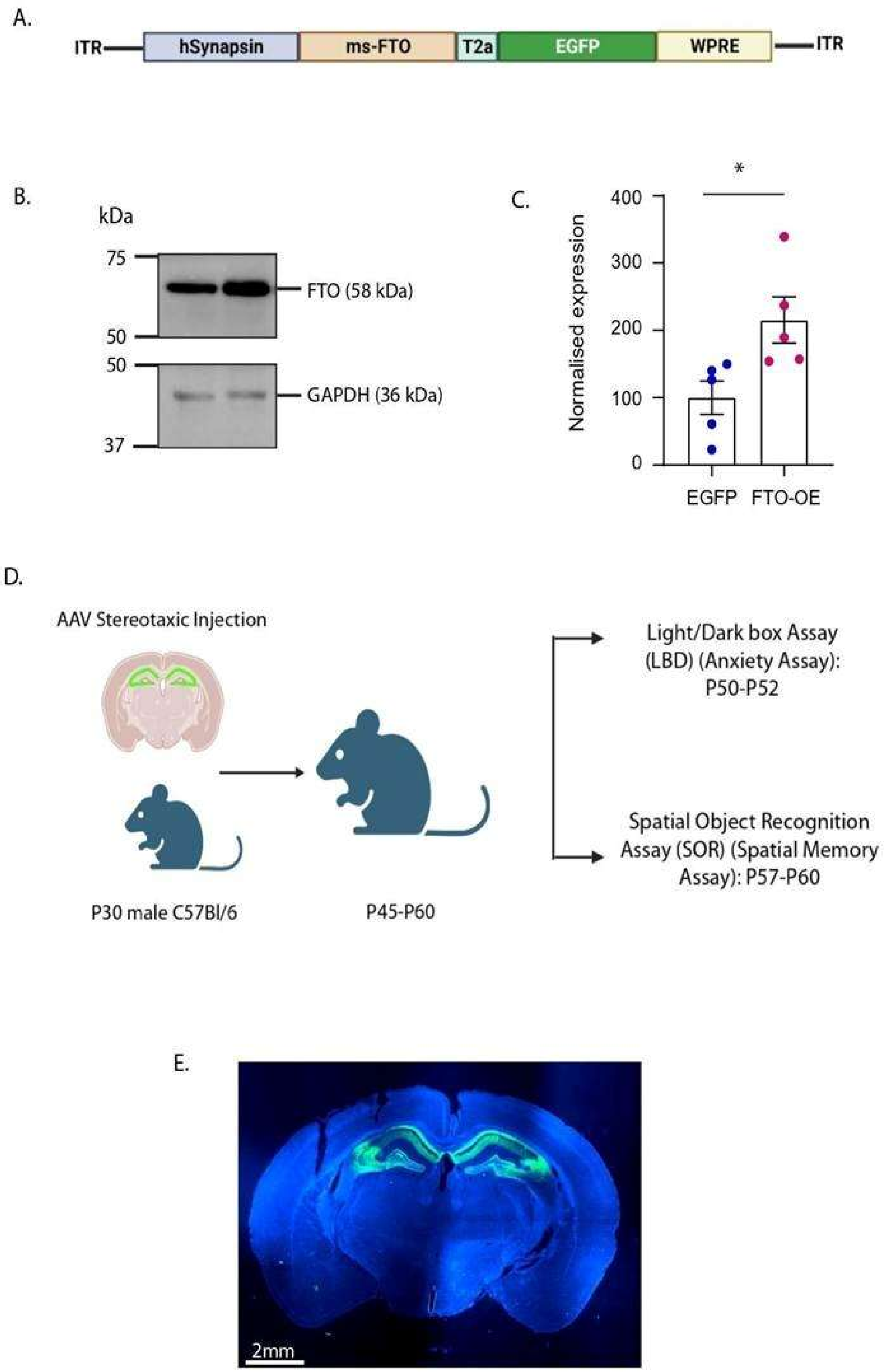


**Supplementary Figure 2. Experimental Design and Validation of FTO Overexpression in the Mouse Brain. (A)** Schematic representation of the AAV vector used for the overexpression of

FTO. **(B)** Immunoblot and **(C)** Quantification for FTO overexpression in HEK293t. Data shown as Mean ± SEM, n=5, *p<0.05. Student's paired t-test. **(D)** Experimental timeline and behavioural assays: AAV inject at postnatal day 30 (P30) into the hippocampus; behaviour assessments between P50 (Anxiety assay: light/dark box assay) and P60 (Spatial memory test: Spatial object recognition test). **(E)** Representative image of coronal brain section showing expression of EGFP, confirming expression of AAV in the hippocampus (P60).

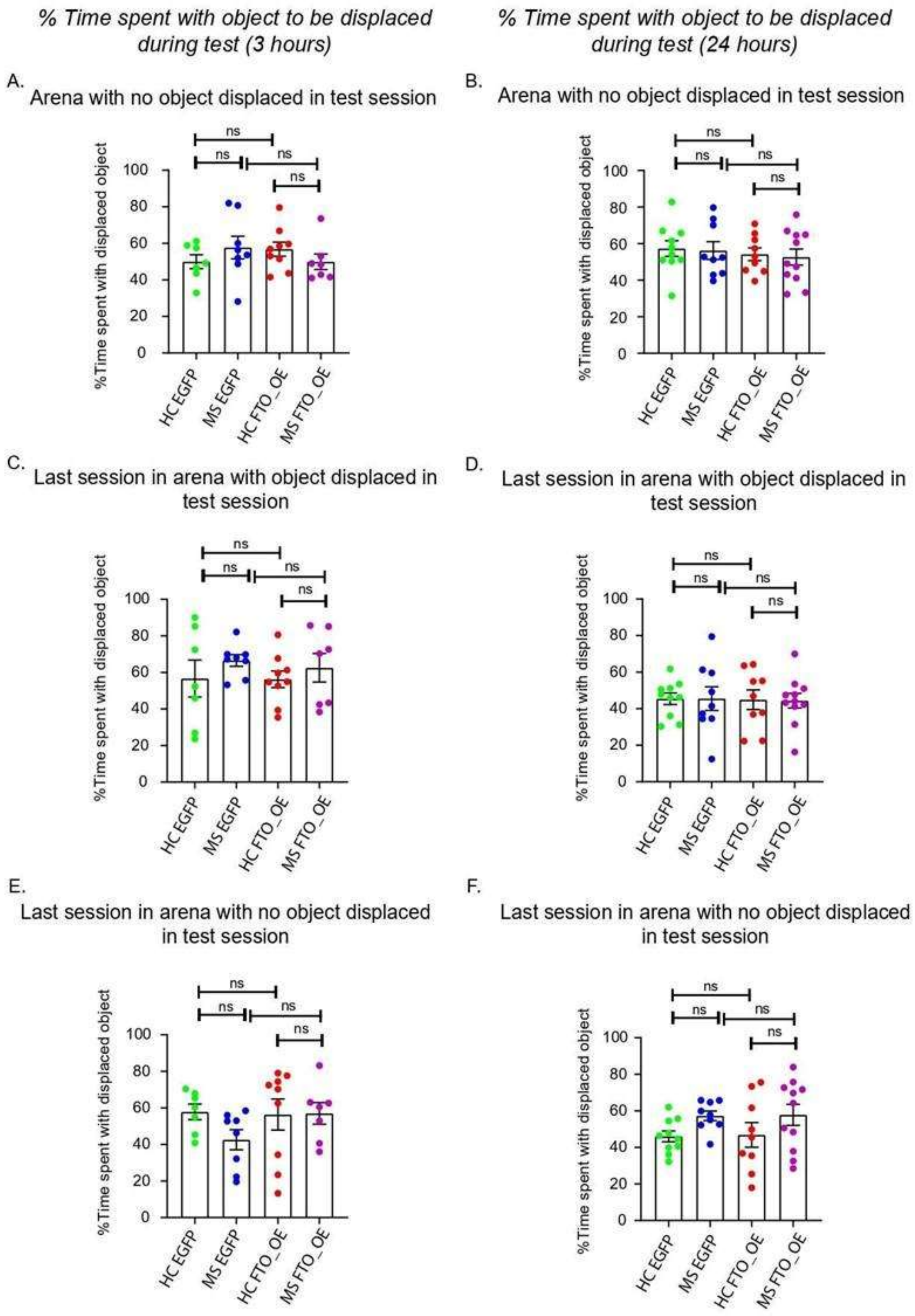


**Supplementary Figure 3. Time spent with object used for SOR shows no object biasness. (A, B)** Percentage time spent with objects in the arena with no object displaced during test session;

**(A)** 3 hours, N=3, n=7-9, and **(B)** 24 hours, N=3, n=9-11, post training. **(C, D)** Percentage time spent in the last training session with objects in the arena with object displaced during test session; **(C)** 3 hours, N=3, n=7-9, and **(D)** 24 hours, N=3, n=9-11, post training. **(E, F)** Percentage time spent in the last training session with objects in the arena with object displaced during test session; **(E)** 3 hours, N=3, n=7-9 and **(F)** 24 hours, N=3, n=9-11, post training. Data shows Mean ± SEM. Two-way ANOVA with Bonferroni's correction, ns = not significant.

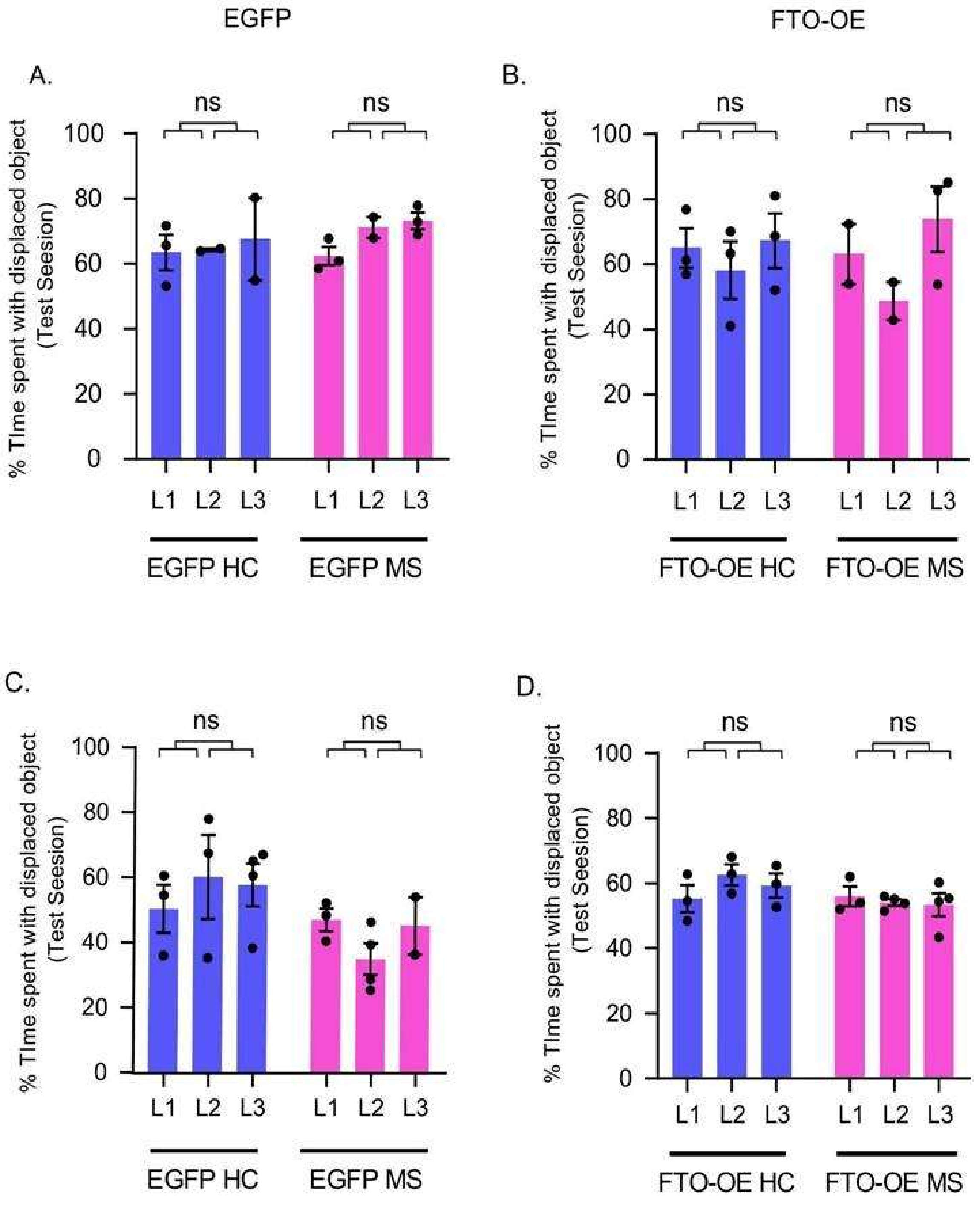


**Supplementary Figure 4**. **Litter effect of animals used for spatial object recognition assay**, Control RNAi, G5C1 RNAi and G5C2 RNAi animals of HC and MS conditions, **(A-B)** test done after 3 hours, N = 3, n=7-9; **(C-D)** test done after 24 hours, N=3, n=9-11. Data shown as mean±SEM. Two-way ANOVA with Bonferroni's correction, ns = not significant.

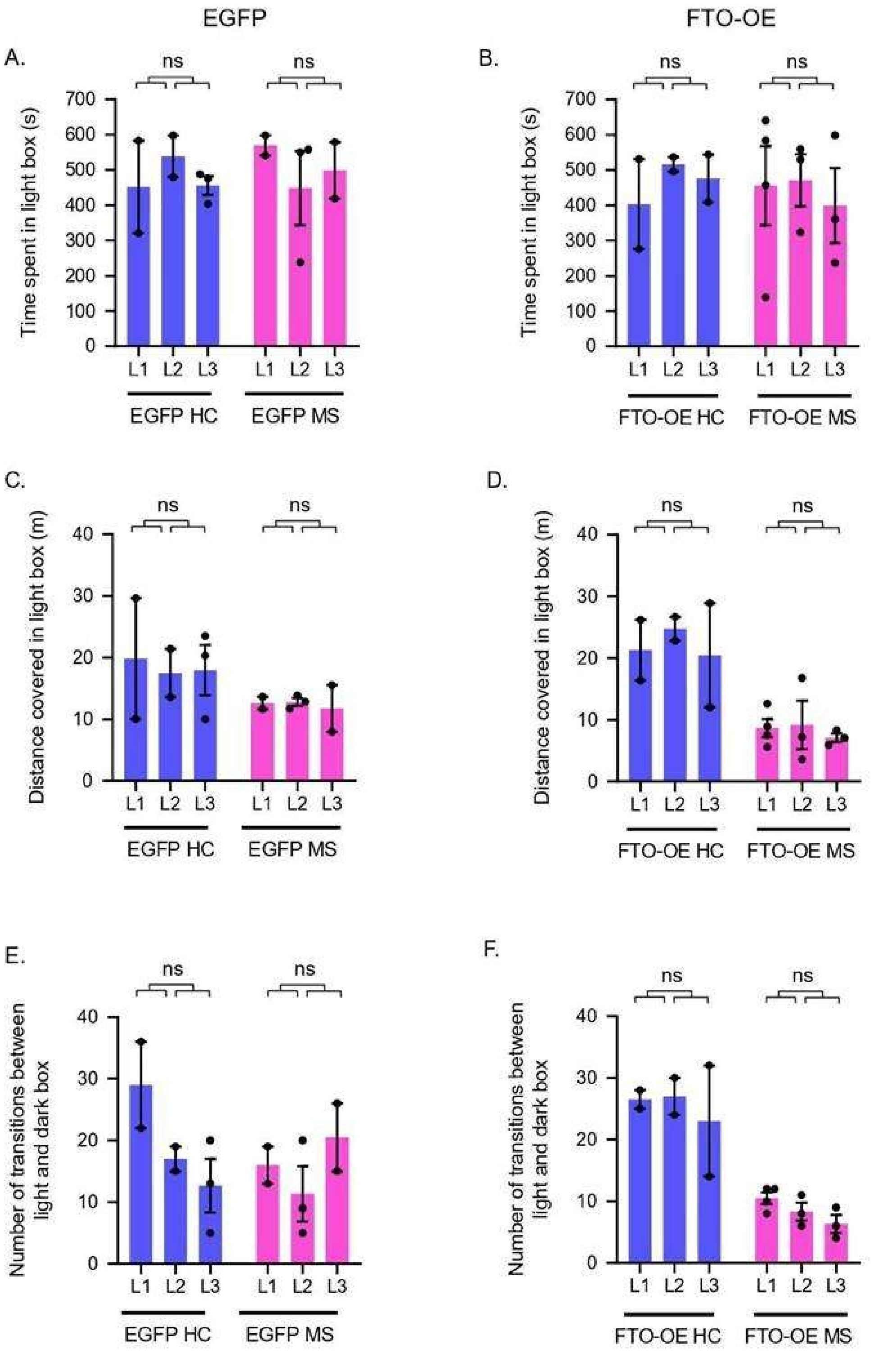


**Supplementary Figure 5. Litter effect amongst animals in all behavioural experiments. (A-C)** Litter effect of animals used for light dark assay for **(A)** Time, **(B)** Distance, **(C)** Transitions in

Control RNAi, G5C1 RNAi and G5C2 RNAi animals of HC and MS conditions. Data shown as Mean ± SEM. Two-way ANOVA with Fisher's LSD, N= 3, n=7-10, ns = not significant.

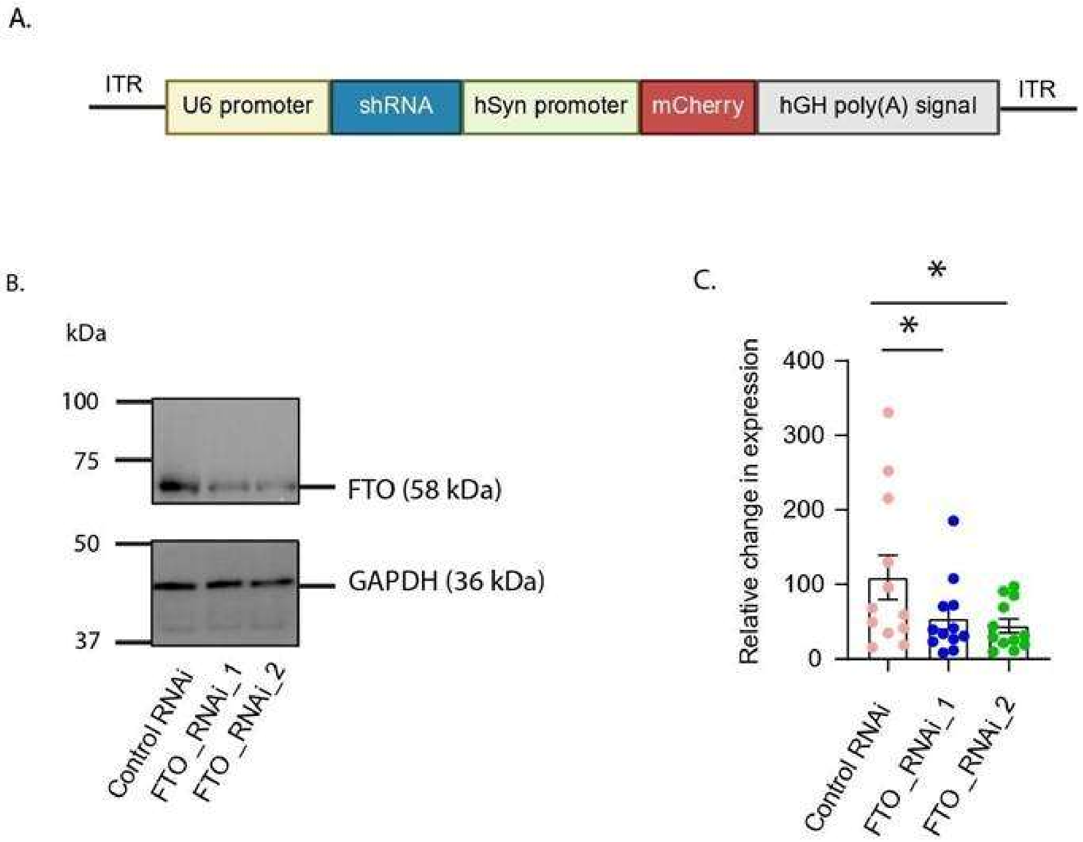


**Supplementary Figure 6. Experimental design and validation of FTO knockdown. (A)** Schematic representation of the AAV vector used for the knockdown of FTO. **(B)** Immunoblot and **(C)** Quantification for FTO knockdown in HEK293t cells on transfection with AAV expressing Control RNAi/FTO RNAi construct 1/ FTO RNAi construct 2. Data shown as Mean ± SEM, n=12, *p<0.05. Student's paired t-test.

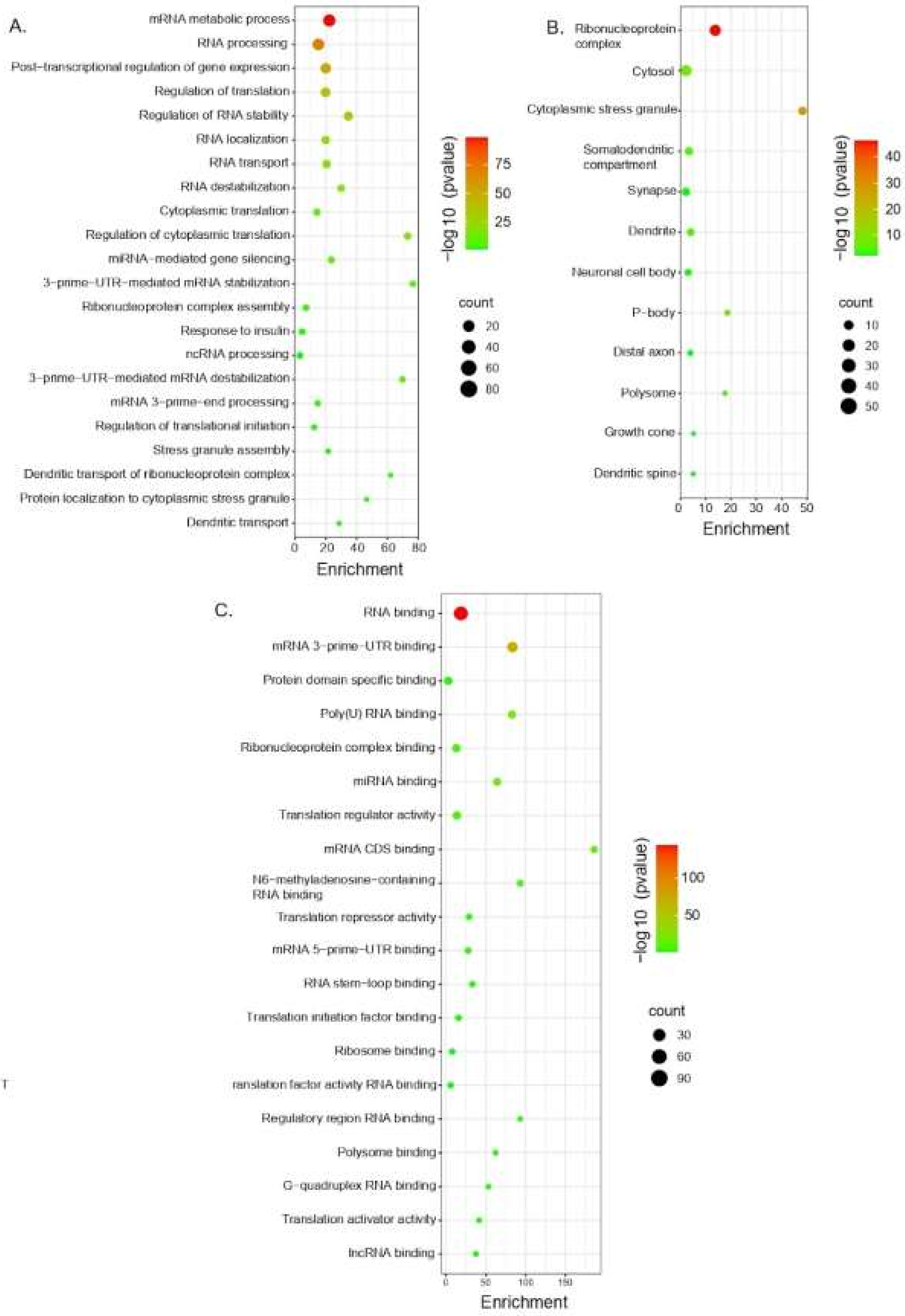
A.
mRNA metabolic process
RNA processing
Post-transcriptional regulation of gene expression
Regulation of translation
Regulation of RNA stability
RNA localization
RNA transport
RNA destabilization
Cytoplasmic translation
Regulation of cytoplasmic translation
miRNA-mediated gene silencing
3-prime-UTR-mediated mRNA stabilization
Ribonucleoprotein complex assembly
Response to insulin
ncRNA processing
3-prime-UTR-mediated mRNA destabilization
mRNA 3-prime-end processing
Regulation of translational initiation
Stress granule assembly
Dendritic transport of ribonucleoprotein complex
Protein localization to cytoplasmic stress granule
Dendritic transport
0 20 40 60 80
Enrichment
-log10 (pvalue)
75
50
25
count
20
40
60
80
B.
Ribonucleoprotein complex
Cytosol
Cytoplasmic stress granule
Somatodendritic compartment
Synapse
Dendrite
Neuronal cell body
P-body
Distal axon
Polysome
Growth cone
Dendritic spine
0 10 20 30 40 50
Enrichment
-log10 (pvalue)
40
30
20
10
count
10
20
30
40
50
C.
RNA binding
mRNA 3-prime-UTR binding
Protein domain specific binding
Poly(U) RNA binding
Ribonucleoprotein complex binding
miRNA binding
Translation regulator activity
mRNA CDS binding
N6-methyladenosine-containing RNA binding
Translation repressor activity
mRNA 5-prime-UTR binding
RNA stem-loop binding
Translation initiation factor binding
Ribosome binding
T
ranslation factor activity RNA binding
Regulatory region RNA binding
Polysome binding
G-quadruplex RNA binding
Translation activator activity
lncRNA binding
0 50 100 150
Enrichment
-log10 (pvalue)
100
50
count
30
60
90

**Supplementary Figure 7. Gene Ontology enrichment and protein-protein interaction network of differentially bound m$^6$A-associated RNA-binding proteins (RBPs). (A)** Biological process, **(B)** Cellular components, and **(C)** Molecular pathways enriched terms, FDR>0.05. **(D)** translation.

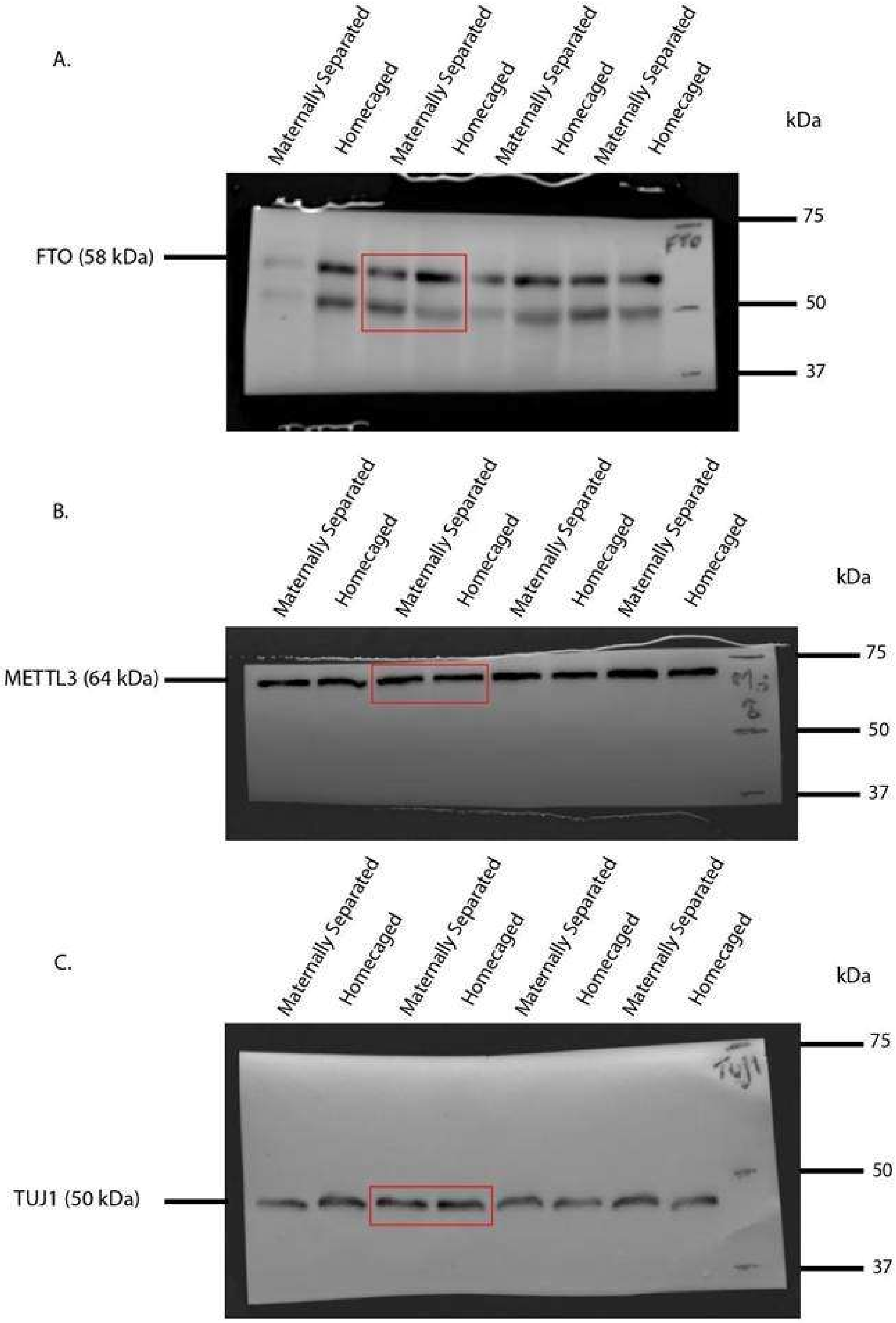


**Supplementary Figure 8.** Full immunoblot showing expression of **(A)** FTO, **(B)** METTL3 and **(C)** Tubulin β1 in males, N=3-4, n=7-11, and **(K)** FTO and METTL3 in P21 males. (See Figure 2 H-J).

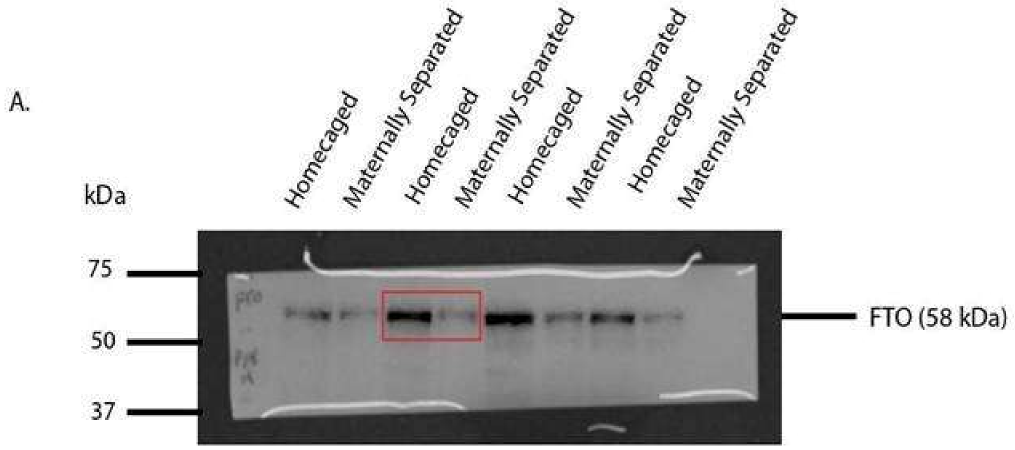


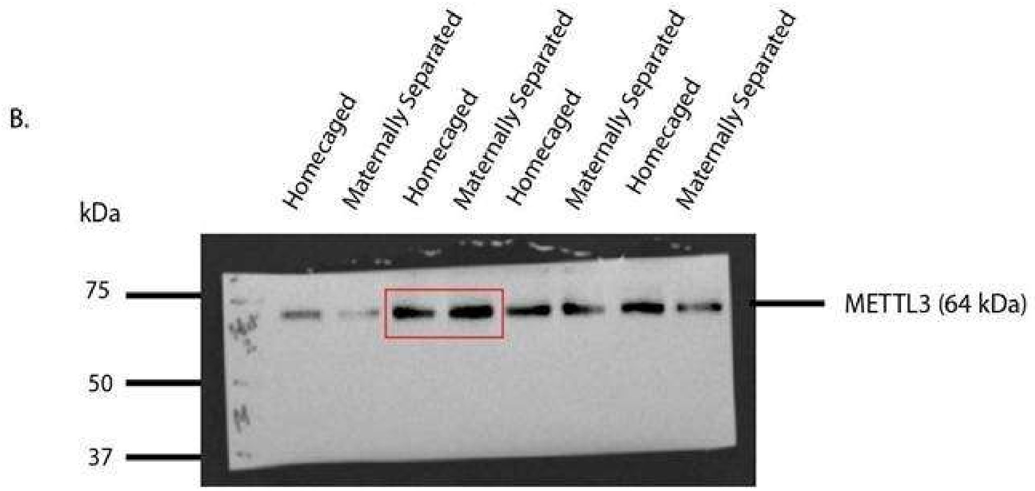


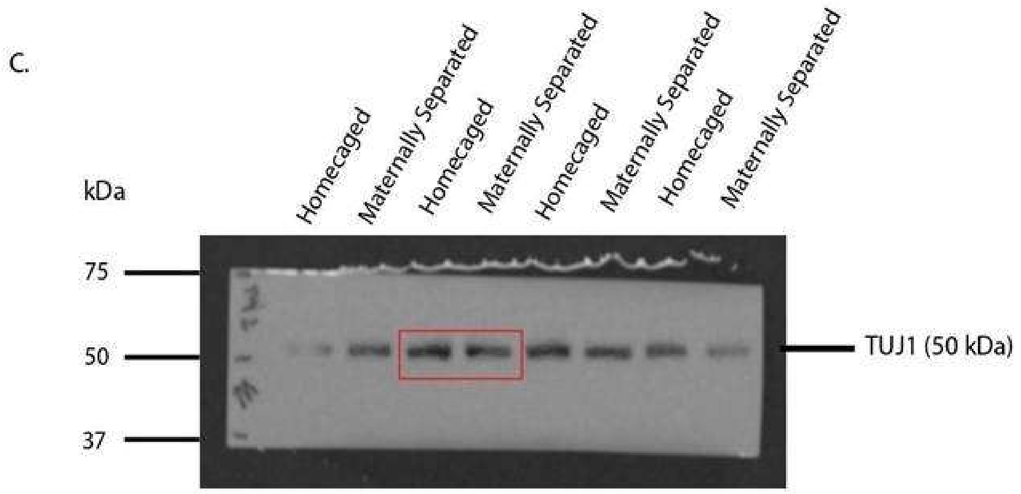


**Supplementary Figure 9.** Full immunoblot showing expression of **(A)** FTO, **(B)** METTL3 and **(C)** Tubulin β1 in males, N=3-4, n=7-11, and **(K)** FTO and METTL3 in P28 males. (See Figure 2 H-J).

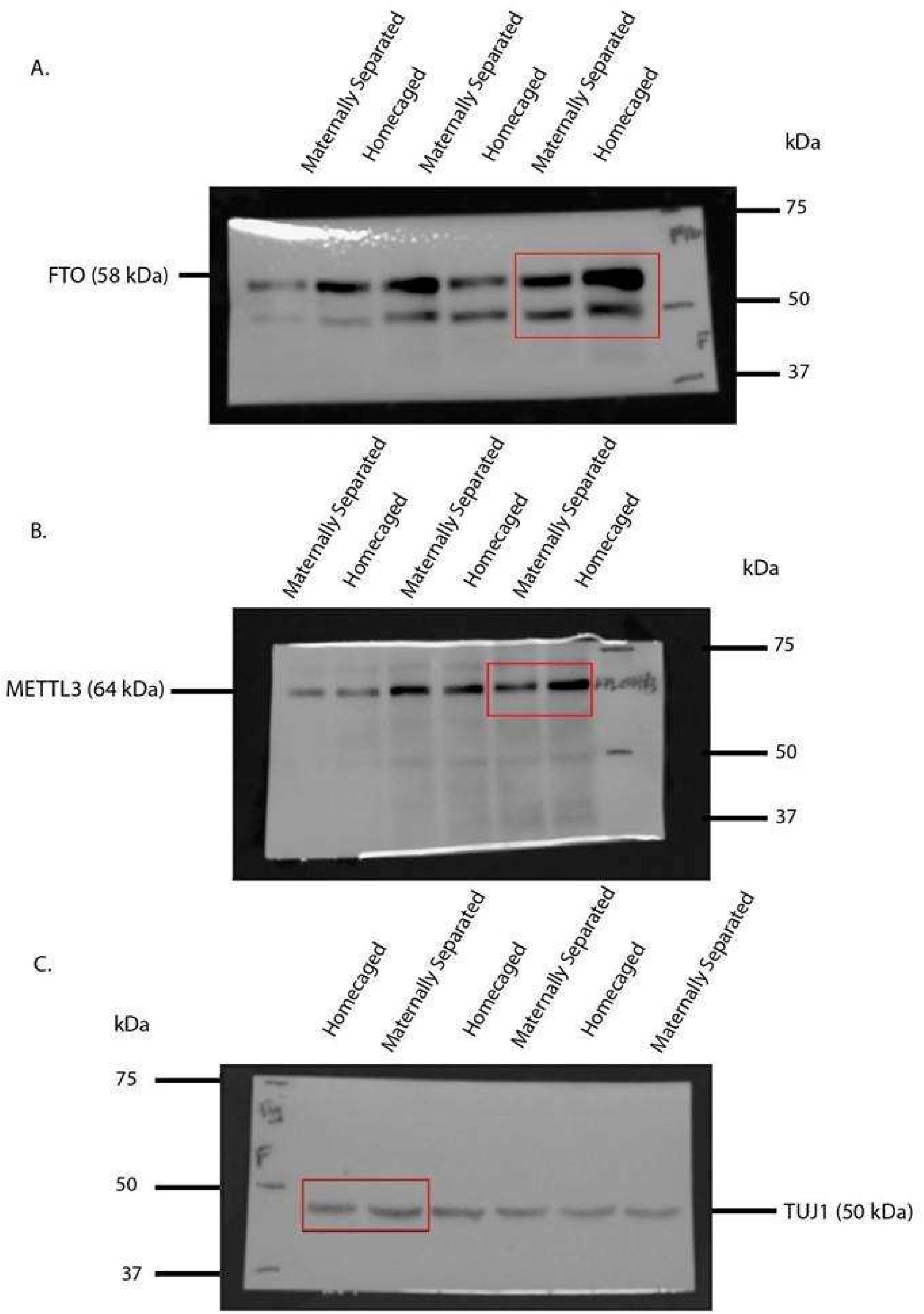


**Supplementary Figure 10.** Full immunoblot showing expression of **(A)** FTO, **(B)** METTL3 and **(C)** Tubulin β1 in P21 females, N=3-4, n=6-7. (See Figure 2 K-M).

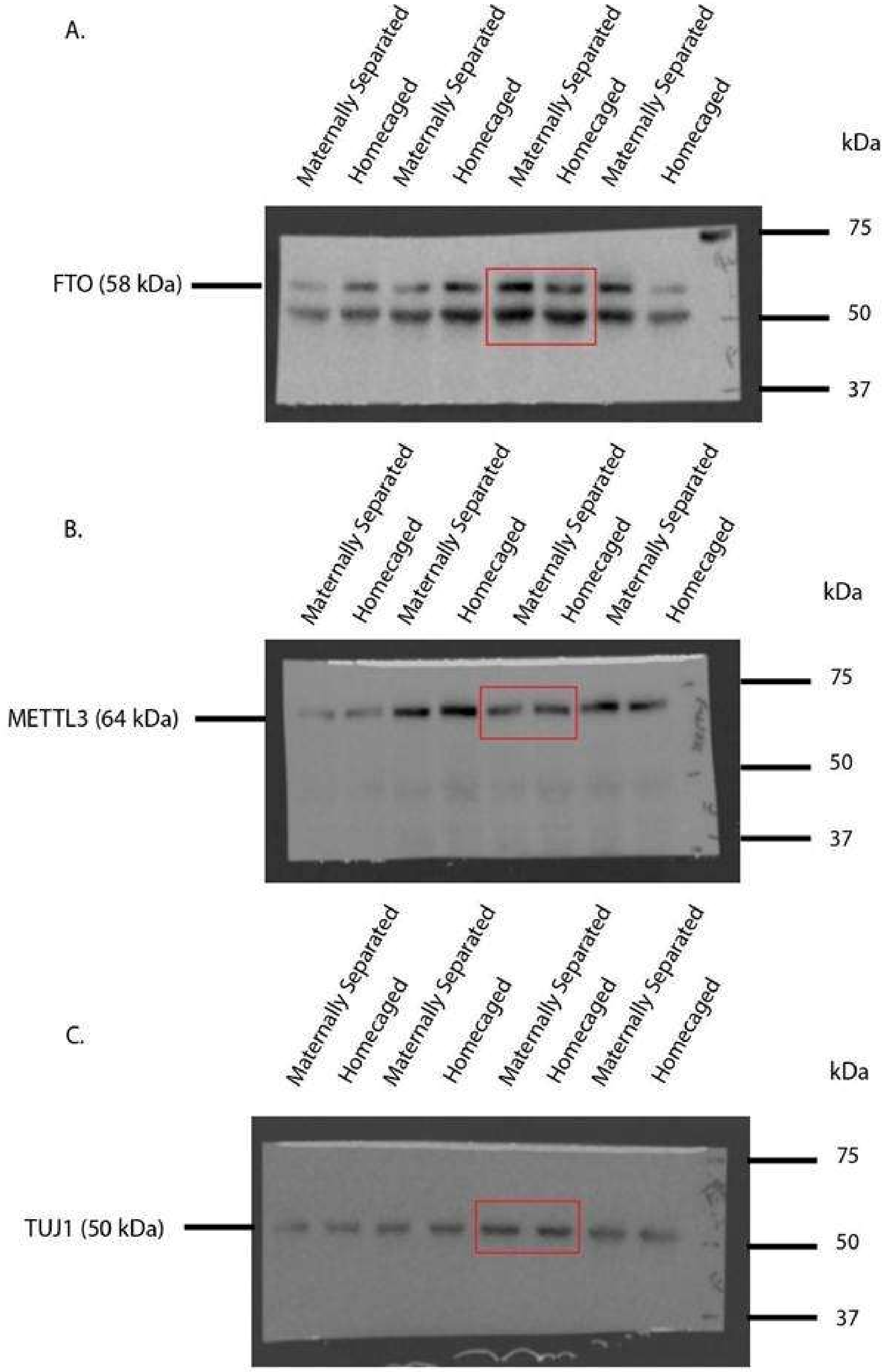


**Supplementary Figure 11.** Full immunoblot showing expression of **(A)** FTO, **(B)** METTL3 and **(C)** Tubulin β1 in P21 females, N=3-4, n=6-7. (See Figure 2 K-M).

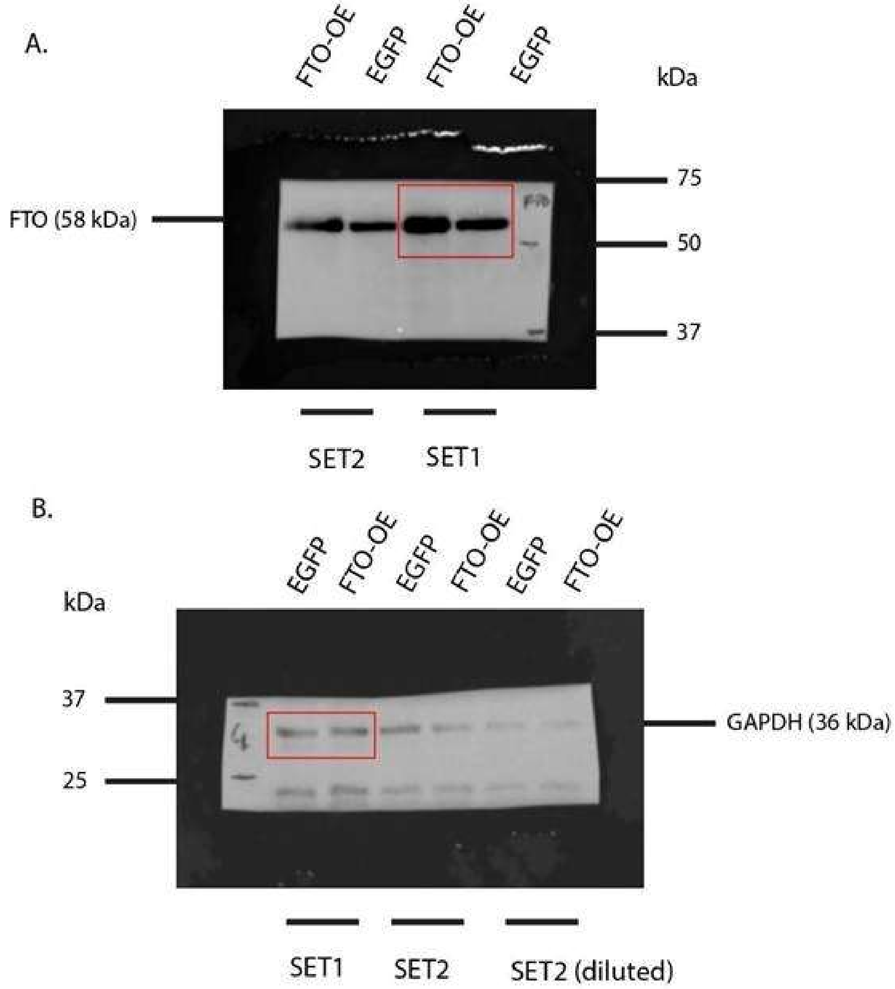


**Supplementary Figure 12.** Full immunoblot showing expression of **(A)** FTO and **(B)** GAPDH. (See Supplementary Figure 1 A-C).

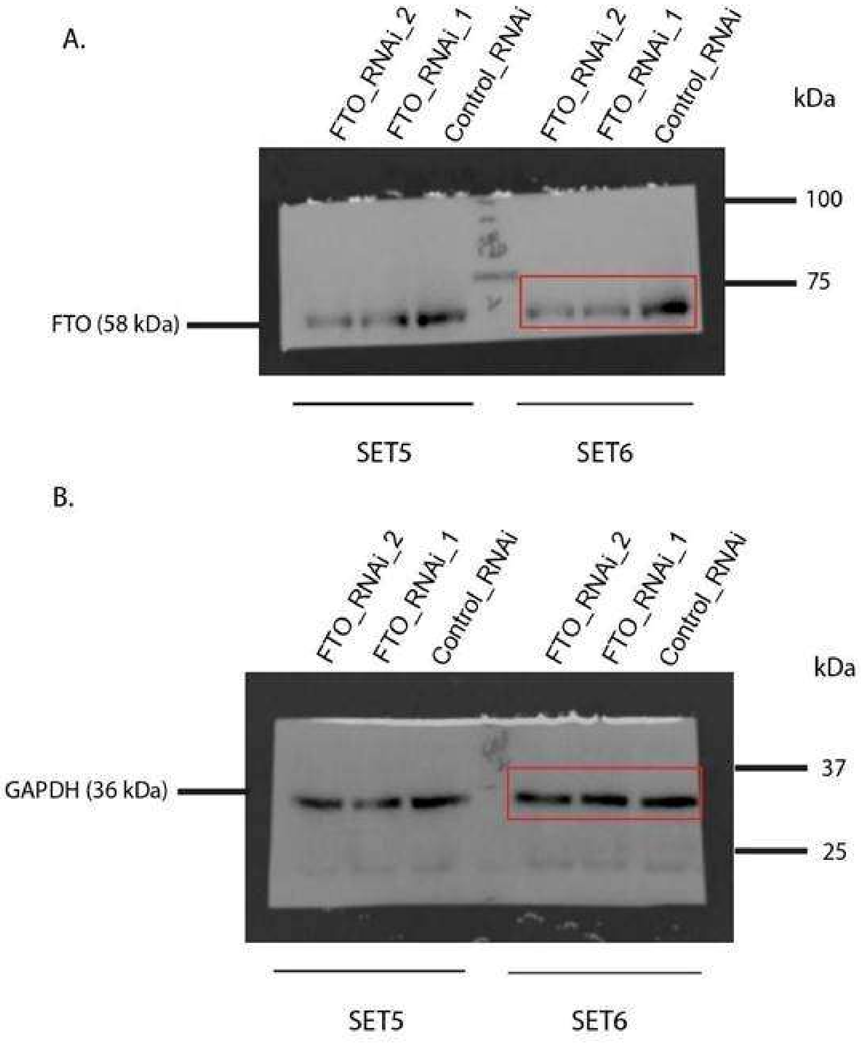


**Supplementary Figure 13.** Full immunoblot showing expression of **(A)** FTO and **(B)** GAPDH. (See Supplementary Figure 5 A-C).
